# The Liver-Enriched Long Non-Coding RNA FAM99A Suppresses Tumorigenesis through Negative Regulation of Protein Synthesis

**DOI:** 10.1101/2025.04.29.651319

**Authors:** Nima Sarfaraz, Ranjit Kaur, Sky Harper, Lilly Oni, Srinivas Somarowthu, Michael J. Bouchard

## Abstract

Primary liver cancer represents a significant global health burden, with limited therapeutic options for advanced disease. Long non-coding RNAs (lncRNAs) are increasingly found to play crucial roles in hepatic biology and disease progression. Here, we identify FAM99A as a highly liver-specific lncRNA that is systematically downregulated across liver malignancies, with reduced expression correlating with poor clinical outcomes. FAM99A exhibits remarkable tissue specificity with minimal expression outside the liver, and its levels rapidly decline during primary hepatocyte dedifferentiation in culture. Through isoform analysis, we establish FAM99A-203 as the predominant transcript in normal liver tissue and observe altered isoform distribution in liver cancers. Functionally, FAM99A overexpression inhibits anchorage-independent growth in liver cancer cell lines. Transcriptomic analysis reveals that FAM99A negatively regulates translation-related pathways in both liver cancer cells and primary hepatocytes. This is corroborated by protein synthesis assays showing that FAM99A overexpression substantially reduces global translation rates. Targeted RNase H-mediated extraction coupled with mass spectrometry identifies multiple components of the translation machinery as direct FAM99A binding partners, including eukaryotic translation initiation factors and RNA helicases involved in ribosome biogenesis. Clinical data analysis demonstrates significant inverse correlations between FAM99A expression and ribosomal protein genes in liver cancer patients. Additionally, hepatitis B virus appears to downregulate FAM99A expression, potentially contributing to its oncogenic properties. Our findings establish FAM99A as a liver-specific translational regulator that exerts tumor-suppressive effects by restraining protein synthesis rates, offering insights into hepatocarcinogenesis and the potential of FAM99A as both a biomarker and agent in new therapeutic avenues.

## INTRODUCTION

Primary liver cancer represents both a formidable challenge in oncology and major global health burden, ranking as the sixth most common cancer worldwide and the third leading cause of cancer-related mortality (1). Hepatocellular carcinoma (HCC), accounting for approximately 90% of cases, develops primarily in patients with chronic liver diseases such as viral hepatitis, alcoholic liver disease, and increasingly, metabolic dysfunction-associated steatotic liver disease (MASLD) (2). Despite advances in surveillance programs for high-risk populations, most patients are diagnosed at intermediate or advanced stages when curative options are limited (3). Current therapeutic approaches, including surgical resection, liver transplantation, locoregional therapies, and systemic treatments, provide modest survival benefits, with median overall survival for advanced HCC remaining under two years (3). The complex molecular landscape of HCC, characterized by substantial heterogeneity and the lack of predominant oncogenic drivers, presents significant obstacles to developing effective targeted therapies (2,4,5). This clinical reality highlights an urgent need to identify novel molecular markers for early detection and targetable pathways for therapeutic intervention. LncRNAs, which are increasingly found to play critical roles in hepatic biology, development, and disease progression, have emerged as promising candidates to address these unmet needs (3,6,7).

FAM99A (Family with Sequence Similarity 99 Member A) is a long intergenic non-coding RNA initially highlighted in transcriptomic studies of liver cancers. Early RNA-seq analyses of HCC tissues identified FAM99A as a differentially expressed lncRNA among others (8). It is predominantly expressed in normal liver (9), though data from expression atlases indicate some detectable levels in placenta, testis, and spleen tissue. In cancer datasets, FAM99A is usually downregulated in liver tumors compared to normal liver. For example, a pan-cancer TCGA survey noted that FAM99A expression is low in liver cancers relative to normal tissue (10). These patterns suggested a potential tumor-suppressive role, prompting focused studies on FAM99A’s function. Interestingly, FAM99A dysregulation has been observed in other disease states, highlighting its broader biological significance beyond hepatic malignancies. One such condition where FAM99A plays a notable role is preeclampsia (PE), a pregnancy complication with significant maternal and fetal health implications. PE is typically characterized by high blood pressure and signs of damage to other organ systems, particularly the kidneys, occurring after 20 weeks of pregnancy. PE involves shallow trophoblast invasion into the maternal spiral arteries, leading to inadequate placental perfusion and release of inflammatory mediators that trigger maternal systemic endothelial dysfunction (11). Several studies feature FAM99A as a regulator of trophoblast cell behavior, where it appears to support normal placental development by enabling trophoblast invasiveness; FAM99A’s aberrant levels in PE correlate with the pathogenic failure of trophoblasts to properly invade the uterine vasculature (12,13).

Multiple patient cohort analyses demonstrate that FAM99A is frequently downregulated in HCC tissues, with low expression correlating with aggressive disease features including higher rates of microvascular invasion and advanced tumor grade (9). Consistently, Kaplan–Meier analyses across multiple datasets reveal that lower FAM99A levels associate with worse overall survival, with multivariate Cox analysis confirming FAM99A as an independent prognostic factor (9). Functional experiments support this clinical correlation; ectopic FAM99A expression in HCC cell lines suppresses proliferation, colony formation, migration, and invasion, while FAM99A knockdown promotes these processes (9). In xenograft models, HCC cells overexpressing FAM99A form significantly smaller tumors, reinforcing its growth-inhibitory role (9). Mechanistically, RNA pull-down coupled to mass spectrometry identified several FAM99A-binding proteins in liver cells, including YBX1, IGF2BP2, HNRNPK, HNRNPL, SRSF5/6, and PCBP (9), many being RNA-binding or ribonucleoprotein factors. Some integrative analyses further suggested FAM99A might participate in hypoxia and metastasis signaling: for instance, one report proposed a hypoxic HDAC1–FAM99A–miR-92a axis promoting HCC metastasis under low oxygen conditions (14). FAM99A may also influence cancer metabolism through regulation of glycolysis. Treatment of HCC cells with icaritin, a plant-derived compound in clinical trials, upregulates FAM99A and suppresses aerobic glycolysis by downregulating glucose transporter GLUT1 (15). FAM99A appears to mediate this effect by binding translation factor EIF4B and reducing synthesis of IL-6 receptor components while simultaneously sponging miR-299-5p to elevate SOCS3, thereby inhibiting the IL6/JAK2/STAT3 pathway and reducing STAT3-driven GLUT1 expression (15).

FAM99B (Family with Sequence Similarity 99 Member B), a paralogous lncRNA located on chromosome 11 within 15-20 kb of FAM99A but on the opposite strand, shares a high degree of sequence similarity with FAM99A. Like FAM99A, FAM99B is highly expressed in normal liver and downregulated in HCC, with low expression correlating with poor clinical outcomes (16,17). Recent studies demonstrate that FAM99B exerts tumor suppression by targeting ribosome biogenesis and protein translation (17). FAM99B physically interacts with DDX21, a nucleolar RNA helicase crucial for rRNA transcription and processing, causing its translocation from the nucleolus to the nucleoplasm and promoting its export to the cytoplasm via XPO1, where it becomes susceptible to proteolytic cleavage (17). This FAM99B-mediated DDX21 degradation impairs ribosome production, limiting the protein synthesis necessary for cancer cell growth (17). Emerging research has also identified a potential role for FAM99B in intercellular communication and therapeutic applications. Xu et al. reported that exosomes derived from human umbilical cord mesenchymal stem cells (MSCs) are enriched in FAM99B and can transfer this lncRNA to HCC cells, inhibiting their malignant phenotype (18). The uptake of exosomal FAM99B induced tumor cell cycle arrest and apoptosis, effectively suppressing HCC growth in vitro (18). This finding highlights exosomes or nanoparticles as potential vehicles for delivering FAM99B to liver tumors (18).

Despite increasing evidence for the tumor-suppressive roles of FAM99A and FAM99B, significant knowledge gaps remain. While DDX21/ribosome interactions have been characterized for FAM99B, detailed mechanistic insights for FAM99A are still emerging. Although sequence homology suggests potential functional overlap between these paralogs, direct experimental evidence confirming shared mechanisms is lacking. Most studies have focused solely on HCC, with minimal expression investigation across other liver cancer subtypes like fibrolamellar carcinoma (FLC) or hepatoblastoma (HPBL). Additionally, while cell line models have provided valuable insights, the behavior of FAM99A in primary human hepatocytes remains uncharacterized. To our knowledge, we are the first group to examine FAM99A function through transcriptomic analysis in both cancer cell lines and primary human hepatocytes, characterize its expression patterns across multiple liver cancer subtypes including FLC and HPBL, perform isoform analyses, and validate its functional significance through protein synthesis regulation in hepatic cells.

## RESULTS

### FAM99A is a Liver-Specific lncRNA Downregulated in Hepatic Malignancies

To identify liver-specific lncRNAs that are dysregulated in HCC, we implemented an integrative bioinformatics approach combining transcriptomic data from multiple sources (Figure 1A). This workflow consisted of three parallel analytical branches: tissue-specific expression profiling using GTEx data, differential expression analysis of HCC versus normal liver tissue using TCGA-LIHC RNA-seq data, and lncRNA annotation using BioMart/GTF resources. From the GTEx dataset, we defined liver-specificity criteria requiring high expression in liver tissue (≥10 TPM) while maintaining minimal expression across all other tissues. Additionally, we imposed a fold-change threshold of ≥5 for liver expression compared to the highest expression value in any non-liver tissue. This stringent filtering identified 18 liver-specific lncRNAs, visualized in the tissue expression heatmap (Figure 1B). The z-score scaled heatmap clearly illustrates the prominent tissue-specificity pattern, with prominent expression (red) exclusively in liver samples while showing minimal to no expression (white/blue) across the remaining tissue types.

**Figure 1:**
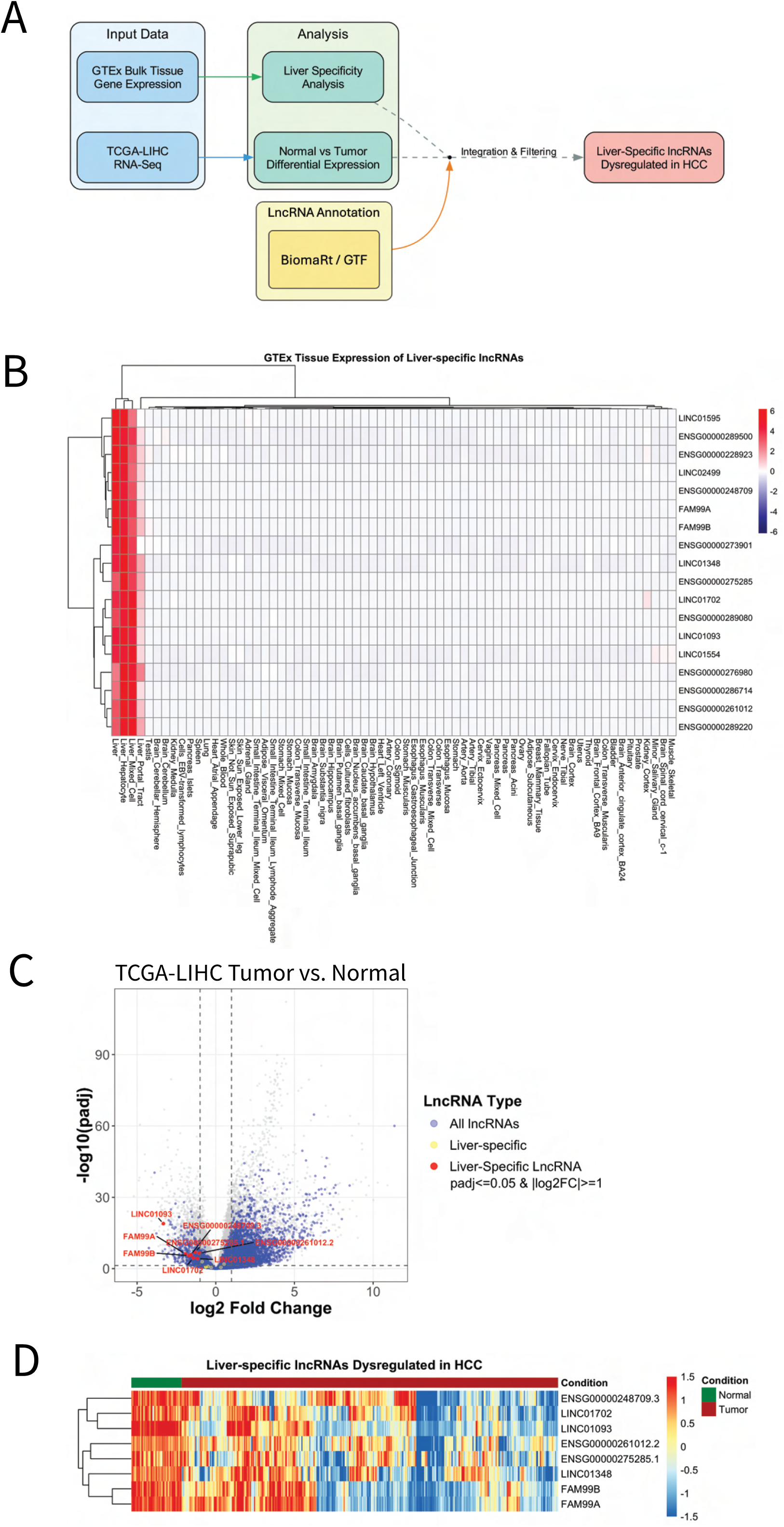
Identification of liver-specific lncRNAs dysregulated in hepatocellular carcinoma (HCC). **(A)** Workflow schematic illustrating the integration of three data sources: TCGA-LIHC RNA-seq for differential expression analysis between tumor and normal liver samples, GTEx bulk tissue gene expression data for tissue-specificity analysis, and lncRNA annotations from BioMart/GTF resources. **(B)** Hierarchical clustering of liver-specific lncRNAs based on expression across 54 human tissues from the GTEx v10 dataset. Heatmap shows row-scaled median TPM values (red = high expression, blue = low expression). Liver-specific lncRNAs (n=18) were defined by expression ≥10 TPM in liver with ≥5-fold higher expression than in any other tissue. **(C)** Volcano plot of differential gene expression in HCC versus normal liver tissue (DESeq2, n=371 HCC tumors, n=50 normal liver samples from TCGA-LIHC). Gray points represent non-significant genes, blue points indicate all annotated lncRNAs), yellow points show liver-specific lncRNAs, and red points highlight liver-specific lncRNAs that are significantly dysregulated in HCC, with gene labels. Vertical and horizontal dashed lines represent log₂FC and significance thresholds, respectively. **(D)** Expression heatmap of the eight liver-specific lncRNAs significantly dysregulated in HCC. Data shown as variance-stabilized transformed (VST) counts from DESeq2, scaled by row. Samples are clustered by condition (green = normal liver, red = HCC tumor).

Concurrently, we performed differential expression analysis comparing HCC tumor samples with matched normal liver tissues from the TCGA-LIHC cohort. After applying statistical thresholds (adjusted p-value < 0.05 and |log₂ fold change| ≥ 1), we identified hundreds of differentially expressed lncRNAs. The volcano plot (Figure 1C) displays the distribution of expression changes. When we intersected these results with our liver-specific lncRNA set, we identified 8 liver-specific lncRNAs that were significantly dysregulated in HCC (red points in Figure 5C), including FAM99A, FAM99B, LINC01702, LINC01093, LINC01348, ENSG00000248709.3, ENSG00000261012.2, and ENSG00000275285.1.

The expression heatmap of these dysregulated liver-specific lncRNAs (Figure 1D) reveals a clear distinction between normal liver (green annotation bar) and HCC (red annotation bar) samples. Normal liver samples consistently showed high expression of these lncRNAs (predominantly red/orange in the heatmap), while HCC samples exhibited frequently reduced expression (predominantly blue/white).

Among these candidates, we selected FAM99A for further characterization based on several compelling features. From the GTEx bulk tissue expression data, FAM99A exhibited the highest median expression level in normal liver tissue (89.04 TPM) among all liver-specific lncRNAs identified, suggesting a potentially critical role in normal liver physiology. Additionally, it displayed substantial downregulation in HCC, ranking as the second most strongly downregulated liver-specific lncRNA (Figure 1C). The existence of its paralog FAM99B, which showed similar expression patterns and downregulation in HCC, further strengthened our hypothesis that FAM99A likely serves an important function in the liver, as evolutionary conservation of this apparent gene duplication event suggested functional significance. These factors collectively positioned FAM99A as a prime candidate for further in-depth functional investigation.

### FAM99A expression is decreased across liver cancer subtypes and correlates with poor prognosis

After identifying FAM99A as a liver-specific lncRNA that is dysregulated in HCC, we conducted focused analyses of its expression patterns and clinical parameters across normal tissues, healthy liver, and various liver malignancies. Analysis of GTEx data confirmed tissue specificity of FAM99A expression (Figure 2A). The lncRNA exhibited extremely high expression in normal liver tissue (median 89.04 TPM), with negligible expression in all other tissues examined. The second highest expression was observed in testes at approximately 3 TPM. This tissue restriction suggests that FAM99A likely functions primarily in hepatocytes and may be involved in liver-specific physiological processes.

**Figure 2:**
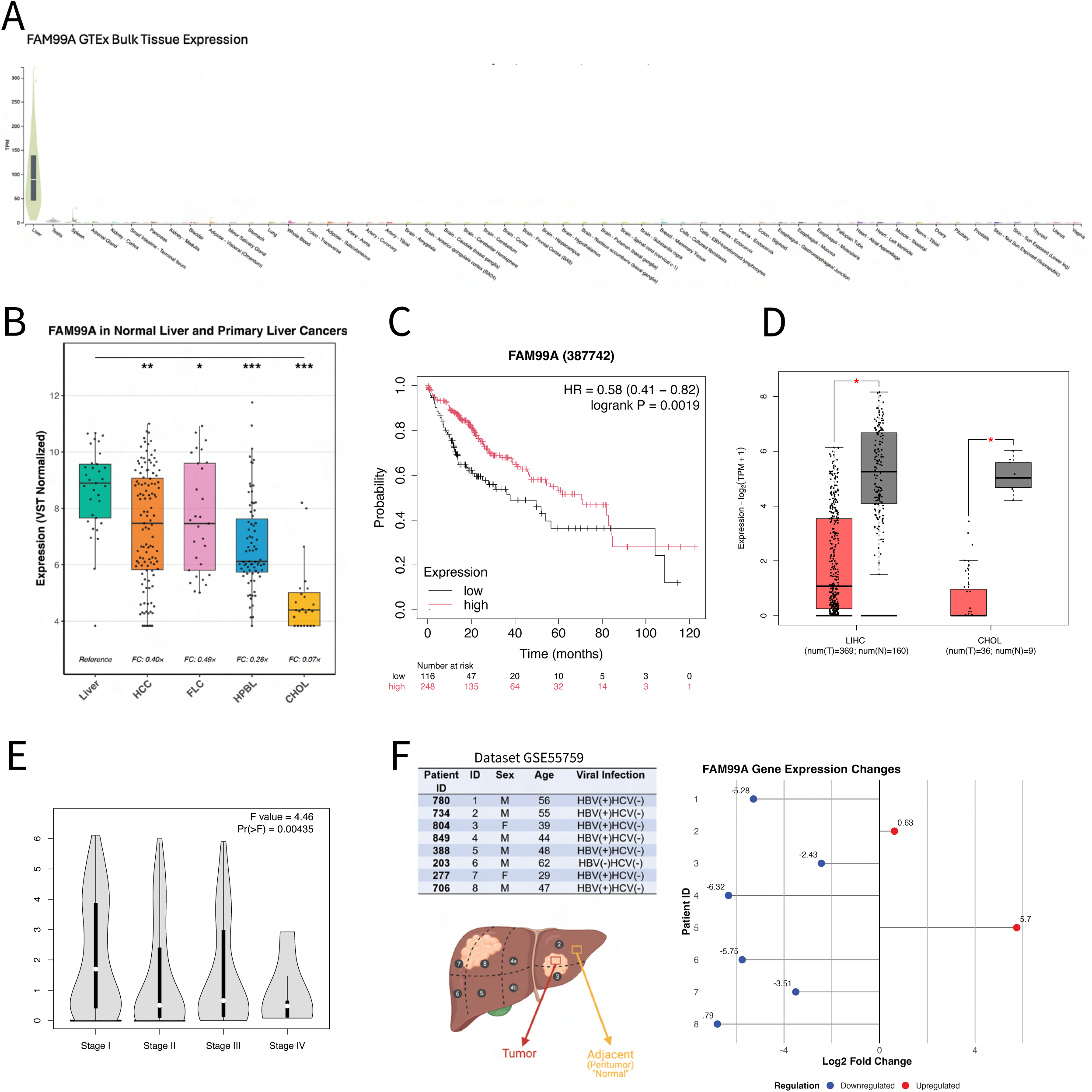
Expression and clinical significance of FAM99A in normal liver and liver malignancies. **(A)** GTEx bulk tissue expression profile of FAM99A across human tissues sorted by median TPM values. **(B)** FAM99A expression across normal liver tissue and primary liver cancer subtypes (VST normalized counts). Statistical significance compared to normal liver: *p < 0.05, **p < 0.01, ***p < 0.001. **(C)** Kaplan-Meier survival analysis of HCC patients stratified by FAM99A expression (HR = 0.58, 95% CI: 0.41-0.82, logrank P = 0.0019). **(D)** FAM99A expression in normal liver versus HCC samples in the TCGA-LIHC cohort (left) and TCGA-CHOL cohort (right), *p < 0.05. **(E)** FAM99A expression across HCC tumor stages I-IV (F value = 4.46, P = 0.00435). **(F)** FAM99A expression in paired HCC tumor and adjacent non-tumor tissues. Analysis of RNA-seq data from GSE55759 dataset showing log2 fold changes in FAM99A expression between tumor and matched adjacent non-tumor tissue from 8 HBV-associated HCC patients. Negative values (blue dots) represent downregulation in tumor tissue compared to adjacent tissue, while positive values (red dots) indicate upregulation. Six of eight patients (75%) exhibited downregulation of FAM99A in tumor tissues

When examining expression across liver cancer subtypes compared to normal liver tissue (Figure 2B), we observed significant downregulation of FAM99A in all liver cancer types. Normal liver samples showed the highest expression levels (VST-normalized mean: 8.66), while HCC samples showed a significant reduction (mean: 7.34, fold change: 0.40×, adjusted p < 0.01). Expression was also reduced in FLC (mean: 7.64, fold change: 0.49×, p < 0.05) and HPBL (mean: 6.72, fold change: 0.26×, p < 0.001). Notably, CHOL samples exhibited the most dramatic reduction (mean: 4.79, fold change: 0.07×, p < 0.001), with expression levels approaching the detection limit. These results indicate that downregulation of FAM99A is a common feature across liver malignancies, though the extent varies by cancer subtype.

The clinical significance of FAM99A expression was assessed using survival analysis in HCC patients (Figure 2C). Kaplan-Meier analysis revealed that patients with higher FAM99A expression demonstrated significantly better overall survival compared to those with lower expression (HR = 0.58, 95% CI: 0.41-0.82, logrank P = 0.0019). This survival advantage was particularly evident beyond 40 months post-diagnosis, suggesting that FAM99A levels may be associated with long-term disease outcomes.

Analysis of the TCGA-LIHC cohort confirmed the differential expression between normal liver and HCC tissues (Figure 2D, left). Normal liver samples showed consistently higher FAM99A expression compared to tumor samples, with small overlap between the groups (p < 0.05). Extending our analysis to TCGA-CHOL, we observed an even more pronounced suppression of FAM99A expression in CHOL tumors compared to normal bile duct tissues (Figure 2D, right), suggesting that loss of FAM99A expression may be a particularly important feature in biliary tract malignancies. Examination of FAM99A expression across HCC tumor stages revealed a significant association with disease progression (F = 4.46, p = 0.00435) (Figure 2E). Expression showed a visible downward trend from Stage I through Stage IV, with the lowest levels observed in Stage IV tumors. This progressive FAM99A loss with advancing disease stage may contribute to the poorer survival observed in patients with lower FAM99A expression.

To further validate our findings from pooled analyses, we examined FAM99A expression in paired tumor and adjacent non-tumor tissues from individual patients using RNA-seq data from the GSE55759 dataset (Figure 2F). This paired analysis approach allows for direct comparison of FAM99A expression changes within the same patient, controlling for individual genetic background and environmental factors. The dataset comprised eight patients (six males and two females) with HCC, with a mean age of 47.5 years (range: 29-62 years). For each patient, samples were obtained from both the tumor and adjacent non-tumor (peritumoral) tissue. In our analysis, six of the eight patients (75%) exhibited significant downregulation of FAM99A in tumor tissue compared to matched adjacent non-tumor tissue (Figure 2F). The predominant pattern of FAM99A downregulation in HCC tumor tissues compared to matched non-tumor tissues from the same patients supports our previous observations in pooled analyses and further strengthens the hypothesis that reduced FAM99A expression may play a role in hepatic carcinogenesis. The observed heterogeneity in expression patterns across patients also highlights the complexity of HCC and suggests that FAM99A dysregulation may vary based on individual tumor characteristics or disease stages.

Collectively, these expression analyses establish FAM99A as a highly liver-specific lncRNA that is systematically downregulated across liver cancer subtypes. The strong correlation between reduced FAM99A expression and poorer survival in HCC patients suggests that this lncRNA may function as a tumor suppressor and could serve as a prognostic biomarker in liver malignancies.

### FAM99A levels rapidly decline during primary hepatocyte culture

We next examined its expression across cultured cell models and primary human hepatocytes. Analysis of RNA-seq data revealed a prominent gradient of FAM99A expression across primary hepatocytes and liver cancer cell lines (Figure 3A). Freshly isolated primary human hepatocytes (PHH_Fresh) exhibited the highest expression levels, serving as a reference point for comparison. PHH cultured under physiological oxygen tension (7% O2, “physioxic”, 36h in culture) maintained substantially higher FAM99A expression compared to those cultured under standard conditions, though still significantly lower than freshly isolated cells (p < 0.001). This finding aligns with previous reports in primary rat hepatocytes under physiological oxygen tension culture conditions can better maintain native hepatocyte physiology (19). Short-term cultured PHH (36h in culture) showed approximately 1.5% of the expression levels observed in fresh cells, while expression in long-term cultured PHH (5 passages and 45 days in culture) dropped further to levels comparable to those seen in liver cancer cell lines. This progressive decline emphasizes the rapid loss of this liver-specific transcript during conventional culture.

**Figure 3:**
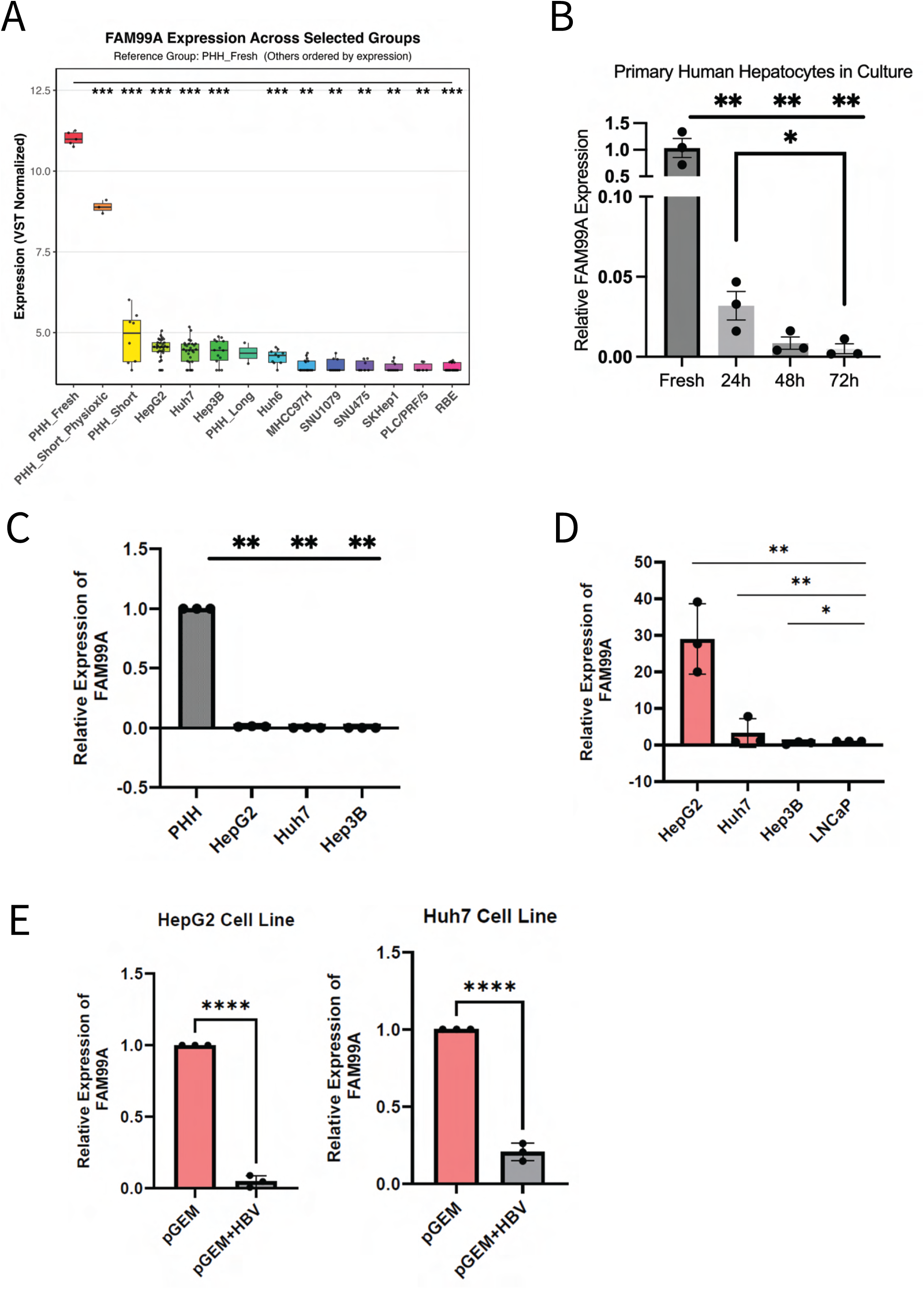
FAM99A expression in primary human hepatocytes and liver cancer cell lines. **(A)** RNA-seq analysis (VST-normalized counts) of FAM99A expression across freshly isolated primary human hepatocytes (PHH_Fresh), primary hepatocytes cultured under different conditions, and liver cancer cell lines. PHH_Short_Physioxic: primary hepatocytes cultured at 7% O2; PHH_Short: primary hepatocytes with maximum one passage; PHH_Long: primary hepatocytes after five passages. All groups show significantly lower expression compared to PHH_Fresh (***p < 0.001, **p < 0.01). **(B)** RT-qPCR analysis of FAM99A expression in primary human hepatocytes from three independent donors during culture at 24, 48, and 72 hours, demonstrating rapid loss of expression. Values normalized to GAPDH and relative to fresh PHH. *p < 0.05, **p < 0.01 versus fresh PHH. **(C)** RT-qPCR comparison of FAM99A expression in fresh PHH versus common liver cancer cell lines. Values normalized to GAPDH. **p < 0.01 versus fresh PHH. **(D)** RT-qPCR comparison of FAM99A expression across liver cancer cell lines and the non-hepatic LNCaP cell line, demonstrating gradient of expression with HepG2 maintaining the highest levels among cancer cell lines. Values normalized to GAPDH and relative to LNCaP. *p < 0.05, **p < 0.01. **(E)** RT-qPCR comparison of FAM99A expression across liver cancer cell lines transfection with a pGEM-HBV plasmid versus control. Values normalized to GAPDH and relative to LNCaP. **p < 0.01, ***p < 0.001.

### Liver cancer cell lines show minimal FAM99A expression

Among liver cancer cell lines, HepG2 cells demonstrated the highest FAM99A expression, followed by Huh7 and Hep3B with progressively lower levels. The poorest expression was observed in cell lines of biliary origin (SNU1079 and RBE), consistent with our previous finding that cholangiocarcinoma tissues exhibit minimal FAM99A expression. Expression in all cell lines was significantly reduced compared to fresh PHH (p < 0.001 for all comparisons).

To validate these transcriptomic findings, we performed RT-qPCR analysis on PHH samples obtained from three independent donors. FAM99A expression declined precipitously during standard culture, dropping to approximately 1% of initial levels by 48 hours (p < 0.01) (Figure 3). This rapid decline parallels the known de-differentiation of hepatocytes in two-dimensional culture (20,21) and correlates with the rise of EMT and YAP signatures observed during early transformation of hepatic cells (20–22), suggesting FAM99A may serve as a sensitive marker of hepatocyte differentiation status.

Direct comparison of FAM99A expression between fresh PHH and commonly used liver cancer cell lines by RT-qPCR confirmed the dramatic difference in expression levels (Figure 3C). HepG2, Huh7, and Hep3B cells all exhibited negligible FAM99A expression compared to fresh PHH (p < 0.01), consistent with our RNA-seq findings. While expression in these cell lines was extremely low relative to primary hepatocytes, a second RT-qPCR experiment comparing only the cell lines to each other revealed that HepG2 cells maintained detectable FAM99A expression at levels approximately 30-fold higher than Huh7 cells (Figure 3D). The non-hepatic LNCaP prostate cancer cell line showed virtually undetectable expression. This gradient of expression across liver-derived cell lines, while still far below physiological levels, suggests that certain liver cancer lines retain a minimal degree of the transcriptional program that drives FAM99A expression. Interestingly, HepG2 or Huh7 cells transfected with a HBV-genome-expressing plasmid demonstrated further loss of FAM99A levels, hinting at a potential role in which HBV may regulate FAM99A levels (Figure 3E), possibly contributing to the viruses’ long-term oncogenic nature.

The marked difference between primary hepatocytes and established cell lines, coupled with the rapid decline in FAM99A expression during culture, highlights the challenges of studying liver-specific lncRNAs in conventional cell models. These findings informed our subsequent approach to functional characterization, where we utilized transient overexpression strategies to restore FAM99A in cell line models.

### FAM99A-203 is the predominant isoform in normal liver tissue

To characterize FAM99A at the transcript level, we first investigated which of the annotated isoforms predominates in normal liver tissue and examined potential alterations in splicing patterns across liver cancer subtypes. The Ensembl database (release 112, May 2024) annotated several potential transcript variants for FAM99A, including FAM99A-201, FAM99A-202, and FAM99A-203 (Figure 4A). These variants differ substantially in their exon inclusion patterns and total transcript length, which could impact the secondary structure and functional domains of the resulting lncRNA.

**Figure 4:**
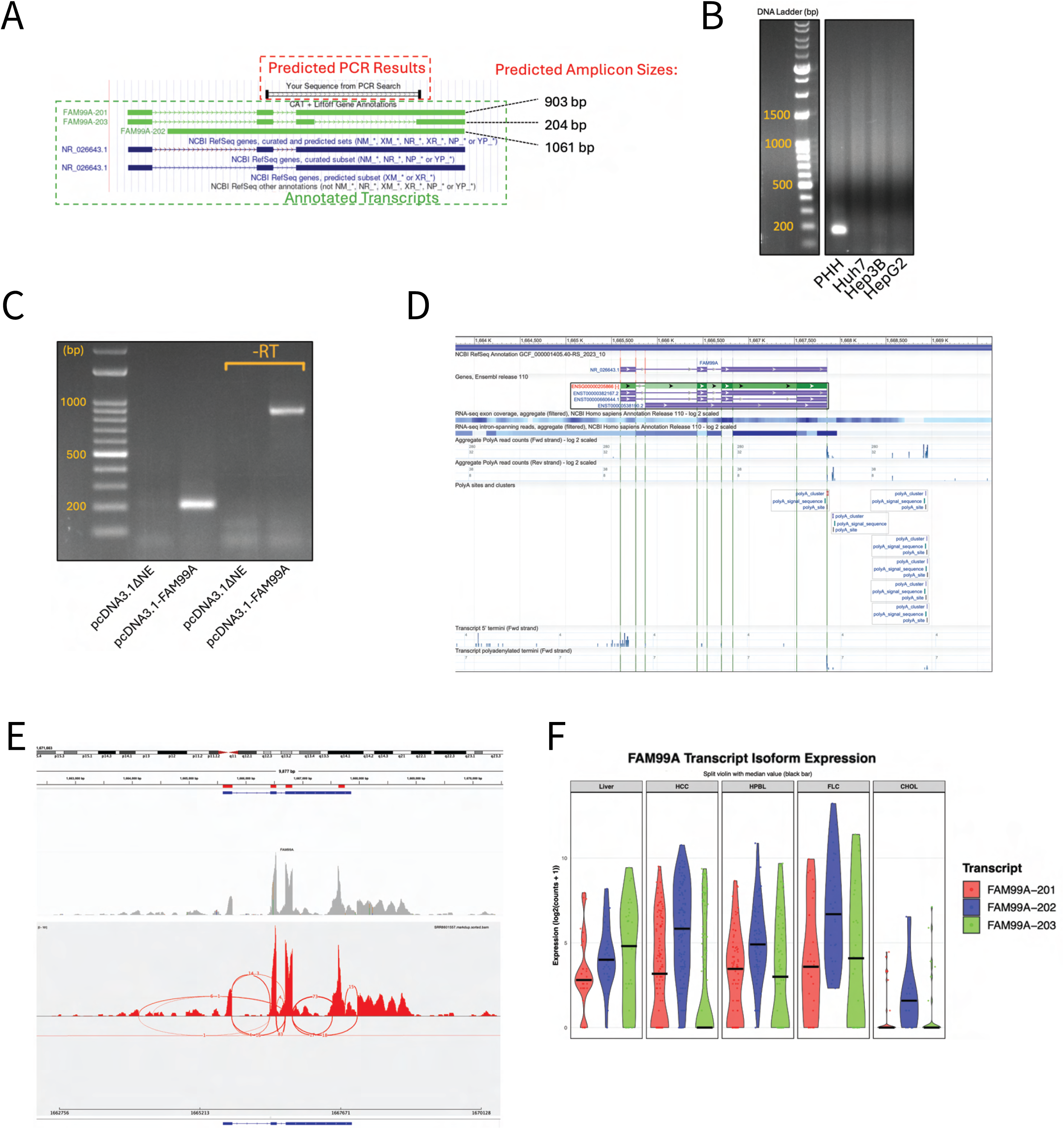
Characterization and validation of FAM99A isoform expression in normal liver and liver cancer. **(A)** In silico PCR analysis showing predicted amplicon sizes for the three major FAM99A transcript variants annotated in Ensembl. Primer design (black bars) targeting shared exons yields diagnostic fragment sizes of 903 bp for FAM99A-201, 204 bp for FAM99A-203, and 1061 bp for FAM99A-202. **(B)** RT-PCR results from freshly isolated primary human hepatocytes (PHH) and liver cancer cell lines (Huh7, Hep3B, HepG2) using isoform-discriminating primers, confirming FAM99A-203 (204 bp) as the predominant transcript in PHH. **(C)** RT-PCR verification of splicing patterns in HepG2 cells transfected with pcDNA3.1-FAM99A (containing FAM99A-201 cDNA). The -RT lanes show residual plasmid DNA (∼900 bp), while the +RT lane demonstrates processing to the FAM99A-203 form (204 bp), indicating preferential splicing to the short isoform even from exogenous constructs. **(D)** Custom track view in UCSC Genome Browser integrating NCBI RefSeq gene annotations with aggregate RNA-seq coverage data. Exon intensity patterns (darker blue) align with FAM99A-203 structure, with additional signals extending beyond annotated transcript boundaries. **(E)** IGV visualization of RNA-seq coverage from a normal liver biopsy sample (SRR8601557), confirming FAM99A-203 as the predominant transcript in normal liver. **(F)** Sashimi plot of the same liver biopsy sample showing splice junction reads (arcs). Thickness of arcs represents junction read counts, with the most abundant splicing events corresponding to the FAM99A-203 isoform. Additional lower-frequency splice junctions connect to regions upstream and downstream of annotated boundaries. **(G)** Quantitative analysis of FAM99A isoform expression across normal liver and liver cancer subtypes (HCC, HPBL, FLC, and CHOL) from RNA-seq data. Split violin plots show not only overall reduction of FAM99A-203 expression in cancer tissues but also a shift in isoform distribution, with increased relative expression of longer FAM99A-202 variant in cancer samples compared to normal liver. Black bars indicate median expression values for each isoform.

We designed a strategic PCR-based approach to discriminate between these isoforms. Primers were positioned at exons shared across variants but flanking differentially spliced regions, yielding predicted amplicon sizes of 903 bp for FAM99A-201, 1061 bp for FAM99A-202, and 204 bp for FAM99A-203 (Figure 4A). RT-PCR analysis of freshly isolated primary human hepatocytes revealed a single dominant band at approximately 200 bp, closely matching the predicted size of the FAM99A-203 variant (Figure 4B). In contrast, the liver cancer cell lines (Huh7, Hep3B, and HepG2) showed substantially reduced amplification of this band, consistent with our previous finding of diminished FAM99A expression in these models.

Our previously synthesized expression plasmid (pcDNA3.1-FAM99A) contained the cDNA sequence corresponding to FAM99A-201. To validate that our expression construct would still correctly reflect the predominant endogenous isoform, we examined the splicing pattern of the transfected FAM99A sequence. Interestingly, when transfected into HepG2 cells, RT-PCR revealed that the exogenous transcript was predominantly processed to yield the 204 bp amplicon matching FAM99A-203 (Figure 4C). The control lanes (-RT) showed a faint band at ∼900 bp, representing residual plasmid DNA, while the +RT lane clearly demonstrated processing to the shorter isoform. This finding suggests that the cellular splicing machinery preferentially produces the FAM99A-203 variant even when provided with the full-length sequence, supporting FAM99A-203 as the predominant functional isoform.

To further validate these observations, we analyzed RNA-seq data using multiple visualization approaches. A custom track view integrating NCBI RefSeq annotations with aggregate RNA-seq coverage data showed the highest exon coverage coinciding with the FAM99A-203 isoform structure (Figure 4D). The exon heatmap intensity clearly demonstrated preferential expression of the regions corresponding to the short isoform. Intriguingly, we observed additional signals extending beyond the annotated 3’ terminus of FAM99A-203, including other potential polyA sites. This suggests the existence of previously unannotated transcript variants. Examination of liver biopsy RNA-seq data in the Integrative Genomics Viewer (IGV) confirmed these findings, with coverage patterns strongly supporting FAM99A-203 as the predominant transcript in normal liver tissue (Figure 4E). The sashimi plot visualization of splice junctions revealed that the majority of splicing events corresponded to those expected for FAM99A-203 (Figure 4F). However, we detected additional splice junctions connecting to downstream regions beyond current annotations, as well as some upstream of the canonical start site, indicating potential novel splice variants deserving future investigation.

Quantitative analysis of isoform expression across primary liver samples provided an intriguing insight into FAM99A transcript dynamics in liver cancers (Figure 4G). Split violin plots depicting isoform-specific expression across liver cancer subtypes revealed that while FAM99A-203 remained the predominant isoform in normal liver, its expression was significantly reduced in HCC, HPBL, FLC, and dramatically diminished in CHOL. Most notably, we observed a relative increase in the proportion of FAM99A-202 (the longer isoform) in cancer samples compared to normal liver. This shift in isoform usage was particularly evident in FLC samples, where FAM99A-202 showed higher median expression than FAM99A-203.

This pattern of altered splicing across liver cancers suggests that FAM99A dysregulation may involve not only reduced overall expression but also a shift toward longer, possibly unprocessed isoforms. Based on the structural differences between these variants, we hypothesize that the shorter FAM99A-203 isoform likely possesses a unique secondary structure that forms the functional domain of this lncRNA. The longer isoforms (FAM99A-201/202) may represent splicing intermediates or variants with altered secondary structures that fail to form this critical functional domain. This finding introduces a new dimension to FAM99A regulation in liver cancer; beyond simple transcriptional silencing, aberrant splicing mechanics may yield non-functional transcripts in a subset of tumors, effectively inactivating FAM99A even when its promoter remains active.

Collectively, these results suggest some un-annotated variants yet to be experimentally validated but establish FAM99A-203 as the predominant isoform in normal liver tissue and demonstrate that liver cancers exhibit both reduced overall expression and altered isoform distribution, potentially contributing to functional inactivation of this lncRNA in malignant states.

### FAM99A overexpression inhibits anchorage-independent growth

Having established FAM99A’s expression patterns across liver cancers and normal tissue, we next investigated its functional impact on cancer cell phenotypes. Since anchorage-independent growth represents a hallmark of malignant transformation, we performed soft agar colony formation assays with Huh7 and Hep3B liver cancer cell lines transfected with either pcDNA3.1-FAM99A or empty vector control. As shown in Figure 5A, Huh7 cells exhibited robust colony formation in the control condition, with numerous distinct colonies visible throughout the field. In contrast, FAM99A-overexpressing Huh7 cells demonstrated markedly reduced colony formation. Quantification of colony density revealed that FAM99A expression significantly suppressed anchorage-independent growth in Huh7 cells, with approximately 50% reduction in colony formation compared to the control (p < 0.05). The effect of FAM99A was also examined in Hep3B cells (Figure 5B). Unlike Huh7 cells, Hep3B cells exhibited lower baseline colony formation efficiency in soft agar, with fewer and smaller colonies observed in the control condition. While FAM99A overexpression appeared to reduce colony formation in Hep3B cells by approximately 30%, this effect did not reach statistical significance. The less pronounced phenotype in Hep3B cells may reflect intrinsic differences in the cell lines’ propensity for anchorage-independent growth or differential sensitivity to FAM99A-mediated growth suppression.

**Figure 5:**
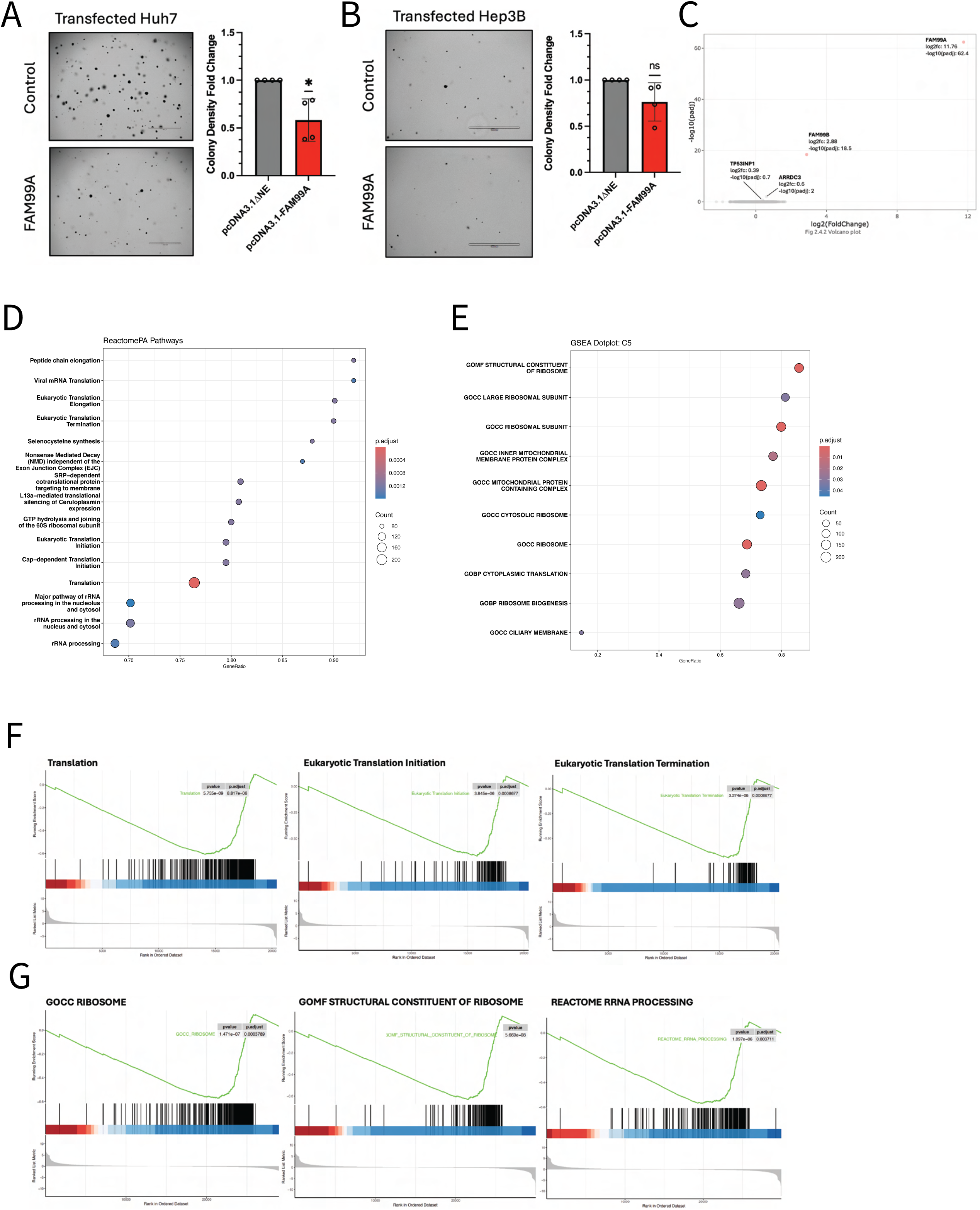
FAM99A suppresses anchorage-independent growth and downregulates translation-related pathways in liver cancer cell lines. **(A)** Left panel: Representative images of soft agar colonies in Huh7 cells transfected with control vector (pcDNA3.1ΔNE) or FAM99A expression construct (pcDNA3.1-FAM99A). Right panel: Quantification of relative colony formation normalized to control. FAM99A overexpression significantly reduced colony formation in Huh7 cells (*p < 0.05, one-sample t-test compared to normalized control value of 1.0, n = 4). **(B)** Left panel: Representative images of soft agar colonies in Hep3B cells transfected with control vector or FAM99A expression construct. Right panel: Quantification of relative colony formation normalized to control. FAM99A overexpression showed a trend toward reduced colony formation in Hep3B cells, though not statistically significant (ns, Wilcoxon signed-rank test compared to normalized control value of 1.0, n = 4). Scale bars = 1000 μm. Error bars represent mean ± SEM. **(C)** Volcano plot showing differential gene expression between FAM99A-overexpressing and control Huh7 cells collected 72 hours post-transfection. FAM99A shows the most significant upregulation (log₂FC: 11.76, -log₁₀(padj): 62.4), with limited changes in other transcripts including TP53INP1 (log₂FC: 0.39) and ARRDC3 (log₂FC: 0.6). **(D)** Reactome pathway analysis of gene set enrichment, revealing significant impacts in translation-related pathways including “Translation,” “Eukaryotic Translation Elongation,” and “Ribosome Biogenesis.” Circle size indicates gene count; color intensity represents adjusted p-value. **(E)** Gene Ontology (GO) analysis using C5 gene sets showing significant enrichment of terms related to ribosome structure and function, rRNA binding, and translation in downregulated genes. **(G)** GSEA enrichment plots for translation and ribosome gene sets, demonstrating coordinated downregulation of translation-related and ribosome biogenesis genes in FAM99A-overexpressing cells. Black vertical bars represent the positions of genes from each gene set in the ranked list; green curves show the running enrichment score.

Regardless, both cell lines showed the same directional trend, with FAM99A expression consistently associated with decreased colony formation. These findings suggest that FAM99A possesses growth-suppressive properties in liver cancer cells, particularly in the Huh7 cell line, which aligns with the hypothesis that FAM99A functions as a tumor suppressor in hepatocellular carcinoma. The more modest effect observed in Hep3B cells might indicate context-dependent activity of FAM99A or may reflect technical limitations of the assay for this particular cell line, which formed fewer colonies overall and might require optimization of parameters such as cell seeding density or incubation time. Together, these results provide functional evidence supporting a growth-inhibitory role for FAM99A in liver cancer cells, consistent with its reduced expression in HCC tissues compared to normal liver.

### Transcriptome analysis reveals FAM99A suppression of translation-related pathways in both Huh7 cells and PHHs

To further investigate the mechanistic impacts of FAM99A in liver cancer cells, we performed RNA-seq analysis on Huh7 cells transiently transfected with either pcDNA3.1-FAM99A or an empty vector control. RNA was collected 72 hours post-transfection to capture stable gene expression changes beyond immediate transfection responses. Differential expression analysis confirmed successful overexpression of FAM99A, with this transcript showing the most significant upregulation (log₂FC: 11.76, -log₁₀(padj): 62.4) in transfected cells (Figure 5C). Interestingly, despite robust FAM99A overexpression, we observed minimal transcriptome-wide changes, suggesting that FAM99A may not primarily function through extensive transcriptional regulation. Among the few significantly altered transcripts were TP53INP1 (log₂FC: 0.39, -log₁₀(padj): 0.7), a known stress-induced p53-regulated gene, and ARRDC3 (log₂FC: 0.6, -log₁₀(padj): 2.0), a tumor suppressor gene involved in β2-adrenergic receptor regulation (23,24).

Given the limited number of high fold-change differentially expressed genes, we employed Gene Set Enrichment Analysis (GSEA) to identify coordinated shifts in pathway activity. This approach revealed significant enrichment of translation-related gene sets among downregulated genes. Reactome pathway analysis (Figure 5D) demonstrated significant negative enrichment in multiple translation-associated processes, including “Translation,” “Eukaryotic Translation Elongation,” “Formation of a pool of free 40S subunits,” and “Ribosome biogenesis.” These findings indicate that FAM99A overexpression induces a coordinated downregulation of translational machinery components. This pattern was further corroborated by Gene Ontology (GO) analysis using the C5 gene set collection (Figure 5E). The most significantly enriched terms among downregulated genes included “Structural Constituent of Ribosome,” “rRNA Binding,” “Cytoplasmic Translation,” and “Ribosome Biogenesis.“

To visualize the specific enrichment patterns, we examined enrichment plots for translation and ribosome related sets (Figures 5F, 5G). Both sets of plots revealed significant negative enrichment scores, with clear rightward skewing of translation-related genes in the ranked dataset. The normalized enrichment scores for “Translation” and “GOCC Ribosome” indicate coordinated downregulation of these pathways. Collectively, these results suggest that while FAM99A does not extensively alter individual gene transcription, it induces a directed negative regulation of translation-related gene programs and ribosome biogenesis. The observed pattern of translational pathway suppression could represent a mechanism by which FAM99A exerts its anti-proliferative effects in liver cancer cells.

To validate and extend our findings from liver cancer Huh7 cells, we investigated the effects of FAM99A overexpression in primary human hepatocytes (PHHs), which represent a more physiologically relevant model of normal liver biology. PHHs from three independent donors were transduced with adenoviral vectors encoding either FAM99A (pAV-FAM99A) or enhanced green fluorescent protein (pAV-GFP) as a control, or left untransduced (Figure 6A). Transduction efficiency was monitored by fluorescence microscopy of pAV-GFP-infected cells, which showed progressive GFP expression from 24 to 72 hours post-transduction, with >80% of cells expressing GFP by the 72-hour timepoint. RNA-sequencing was performed on samples collected 72 hours post-transduction to assess transcriptome-wide changes induced by FAM99A overexpression. To identify FAM99A-specific effects while controlling for both adenoviral transduction and GFP expression, we implemented a carefully designed analytical approach. Both pAV-GFP and pAV-FAM99A samples underwent the same adenoviral transduction process, which inherently controlled for general transduction effects in the direct comparison between these groups. Additionally, we performed a separate analysis comparing untransduced versus pAV-GFP samples to identify pathways affected by GFP expression itself. For pathway analysis, we first conducted Gene Set Enrichment Analysis (GSEA) on both comparisons: untransduced versus pAV-GFP (identifying GFP-associated pathways) and pAV-GFP versus pAV-FAM99A (identifying potential FAM99A effects). To isolate truly FAM99A-specific pathways, we then filtered out from the pAV-GFP versus pAV-FAM99A results any pathways that were also significantly altered in the untransduced versus pAV-GFP comparison. This subtraction approach (pAV-EGFP versus pAV-FAM99A) - (untransduced versus pAV-GFP) yielded pathways uniquely modulated by FAM99A, unconfounded by either adenoviral transduction or GFP expression (Figure 6B, 6C).

**Figure 6:**
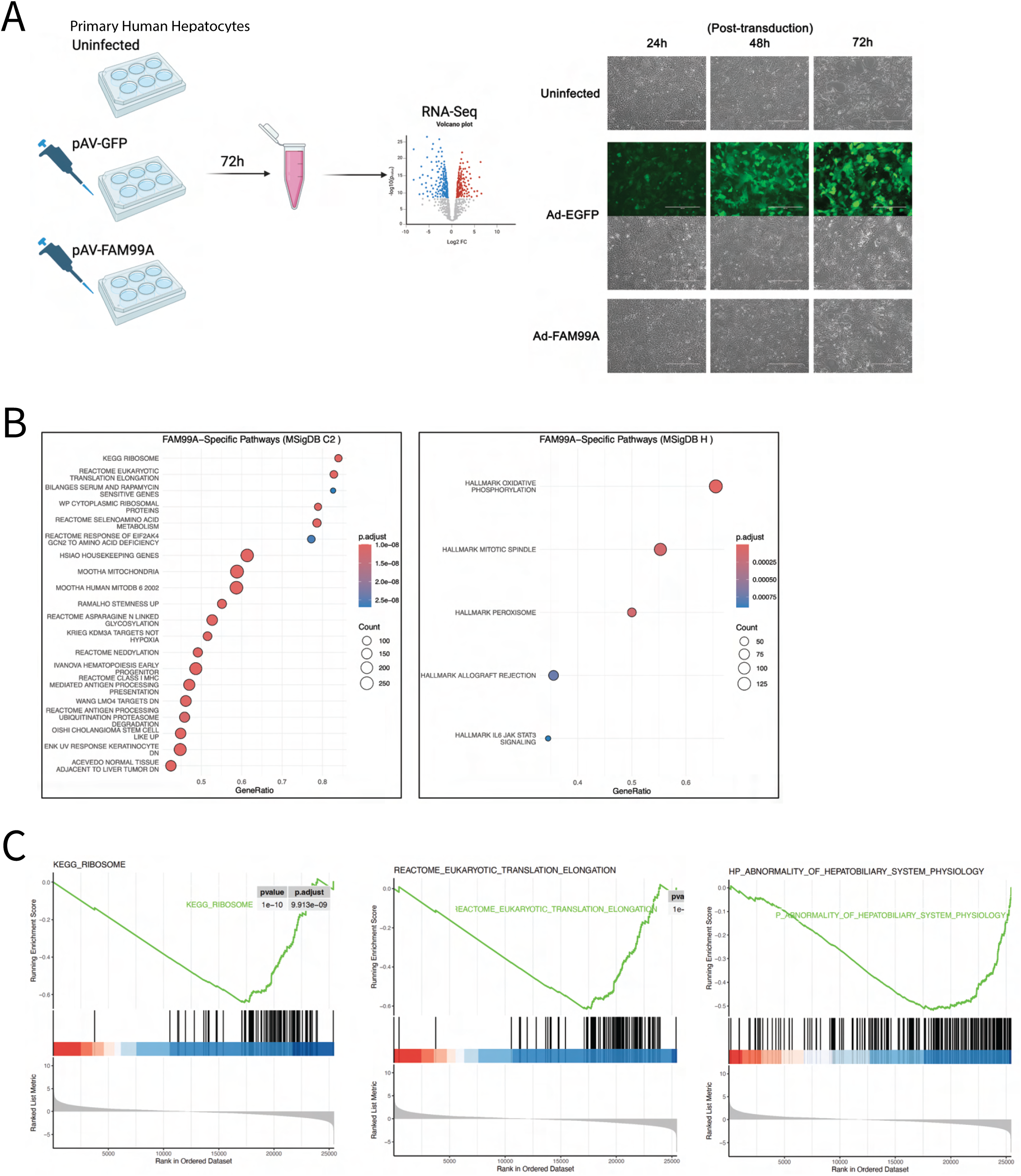
Transcriptomic analysis of FAM99A overexpression in primary human hepatocytes reveals downregulation of translation and ribosome related pathways. **(A)** Left panel: Experimental workflow for adenoviral transduction of primary human hepatocytes. PHHs from three independent donors were either left untransduced or transduced with adenovirus expressing EGFP (Ad-EGFP) or FAM99A (Ad-FAM99A). Cells were collected 72 hours post-transduction for RNA-sequencing analysis. Right panel: Representative brightfield and fluorescence images of PHHs at 24, 48, and 72 hours post-transduction showing adenoviral transduction efficiency. Scale bars = 1000 μm. **(B)** Dotplot visualization of FAM99A-specific pathways identified by Gene Set Enrichment Analysis (GSEA) after filtering out pathways affected by GFP expression. Left panel shows MSigDB C2 collection pathways, highlighting ribosome, translation, and mitochondria-related gene sets. Right panel shows MSigDB Hallmark collection pathways, including oxidative phosphorylation and IL6/JAK/STAT3 signaling. Dot size represents gene count, and color intensity indicates statistical significance. **(C)** Enrichment plots for three representative translation-related pathways downregulated in FAM99A-overexpressing hepatocytes: KEGG_RIBOSOME (left), REACTOME_EUKARYOTIC_TRANSLATION_ELONGATION (center), and HP_ABNORMALITY_OF_HEPATOBILARY_SYSTEM_PHYSIOLOGY (right). All pathways exhibit strong negative enrichment scores with highly significant p-values (p = 1e-10) and adjusted p-values.

In contrast to our observations in Huh7 cells, PHHs exhibited a more robust transcriptional response to FAM99A overexpression, with a larger number of significantly altered genes. The FAM99A-specific pathways identified through this stringent filtering approach showed remarkable concordance with those identified in the cancer cell line model; the FAM99A-specific GSEA revealed pronounced negative enrichment of translation and ribosome biogenesis related pathways (Figure 6B, 6C). The most significantly affected pathways included “KEGG_RIBOSOME” (p-value = 1e-10, adjusted p-value = 9.91e-09), “REACTOME_EUKARYOTIC_TRANSLATION_ELONGATION” (p-value = 1e-10), and “GOCC_CYTOSOLIC_RIBOSOME” (p-value = 1e-10, adjusted p-value = 1.94e-08), all displaying strong negative enrichment scores (Figure 6C). This consistent downregulation of ribosomal and translation-related genes across multiple pathway databases strongly supports FAM99A’s role in regulating translational processes in primary hepatocytes.

Interestingly, analysis of FAM99A-specific responses using MSigDB hallmark gene sets (Figure 6B, right panel) revealed additional pathways uniquely modulated by FAM99A, including “HALLMARK_OXIDATIVE_PHOSPHORYLATION” and “HALLMARK_IL6_JAK_STAT3_SIGNALING.” The identification of IL6/JAK/STAT3 signaling is particularly noteworthy as it aligns with previous reports linking FAM99A to this pathway in the context of icaritin treatment (25), as described in the introduction. The enrichment of oxidative phosphorylation pathways suggests that FAM99A may also influence mitochondrial function and cellular energetics in hepatocytes, potentially connecting translational regulation to metabolic processes.

Additional FAM99A-specific pathways identified from the MSigDB C2 collection (Figure 6B, left panel) included multiple mitochondria-related gene sets (e.g., “MOOTHA_MITOCHONDRIA”), further supporting a potential role in regulating cellular metabolism. We also observed enrichment of gene sets related to asparagine N-linked glycosylation, redox biology, and response to amino acid deficiency, suggesting that FAM99A may coordinate cellular responses to metabolic stress conditions.

Taken together, these findings in primary human hepatocytes corroborate and extend our observations in Huh7 cells, establishing translational regulation as a primary function of FAM99A in both normal and malignant liver cells. The consistent suppression of translation-related pathways across cell models suggests that FAM99A acts as a negative regulator of protein synthesis in hepatocytes. The additional connections to oxidative phosphorylation and JAK/STAT signaling highlight potential mechanisms through which FAM99A might integrate translational control with cellular metabolism and stress responses in the liver.

### FAM99A reduces global protein synthesis rates

To determine whether the transcriptomic findings of translation pathway downregulation translated to functional effects, we assessed the impact of FAM99A overexpression on global protein synthesis rates. We employed a fluorescence-based protein synthesis assay utilizing O-propargyl-puromycin (OPP) incorporation, which allows quantitative assessment of active translation. OPP is a puromycin analog that incorporates into nascent polypeptide chains during translation elongation, effectively labeling actively translating ribosomes. The alkyne group in OPP permits subsequent detection via copper-catalyzed click chemistry with a fluorescent azide.

Hep3B and Huh7 cells were transiently transfected with either pcDNA3.1-FAM99A or empty vector control, and 24 hours post-transfection, cells were incubated with OPP, fixed, stained with 5-FAM-azide, and fluorescence was quantified. As shown in Figures 7A and 7B, FAM99A overexpression induced a substantial reduction in protein synthesis rates in both cell lines. In Hep3B cells, FAM99A expression decreased OPP incorporation by approximately 65% compared to control cells (p < 0.05). Similarly, in Huh7 cells, FAM99A expression reduced protein synthesis rates by approximately 70% relative to control (p < 0.05). Fluorescence microscopy of the labeled cells confirmed these findings, with FAM99A-expressing cells displaying markedly reduced fluorescence intensity compared to control cells.

**Figure 7:**
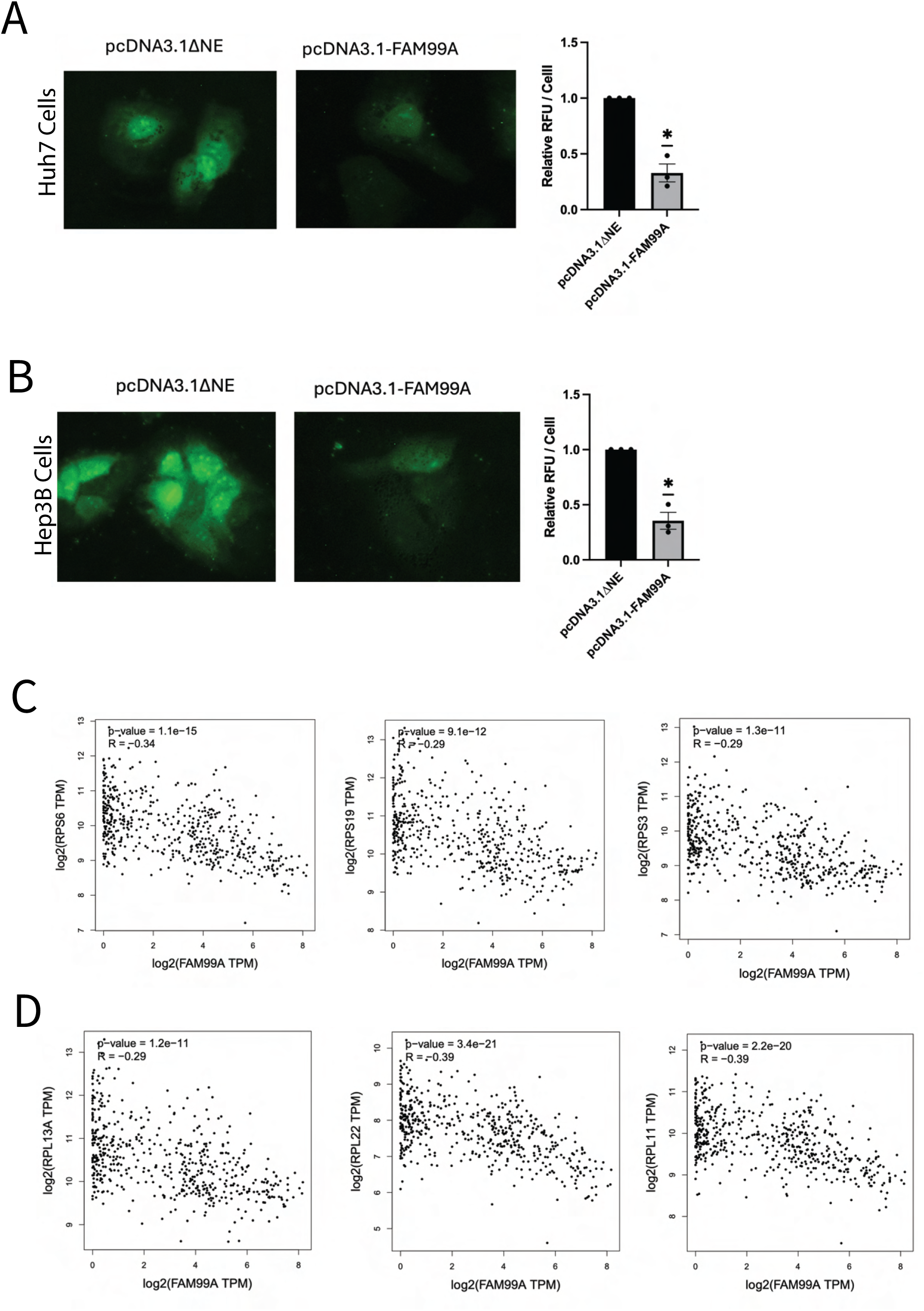
FAM99A reduces global protein synthesis rates in liver cancer cells. **(A)** FAM99A overexpression significantly reduces protein synthesis rates in Huh7 liver cancer cells. Left panels: Representative fluorescence microscopy images of cells transfected with either control plasmid (pcDNA3.1ΔNE) or FAM99A expression construct (pcDNA3.1-FAM99A) and labeled with OPP followed by FAM-based click chemistry. Right panels: Quantification of relative fluorescence intensity measured by plate reader, normalized per cell and to control. Data are presented as mean ± SEM from three independent experiments. *p < 0.05, one-sample t-test compared to a hypothetical mean of 1.0. **(B)** FAM99A overexpression significantly reduces protein synthesis rates in Hep3B liver cancer cells. Left panels: Representative fluorescence microscopy images of cells transfected with either control plasmid (pcDNA3.1ΔNE) or FAM99A expression construct (pcDNA3.1-FAM99A) and labeled with OPP followed by FAM-based click chemistry. Right panels: Quantification of relative fluorescence intensity measured by plate reader, normalized per cell and to control. Data are presented as mean ± SEM from three independent experiments. *p < 0.05, one-sample t-test compared to a hypothetical mean of 1.0. **(C)** Pearson correlation analysis of FAM99A expression with 40S ribosomal protein genes across liver cancer samples. Scatter plots showing significant negative correlations between FAM99A expression and key 40S ribosomal subunit components including RPS6 (R = −0.34, p = 1.1e-15), RPS19 (R = −0.29, p = 9.1e-12), and RPS3 (R = −0.29, p = 1.3e-11) in TCGA liver cancer data. Each point represents an individual patient sample, with regression lines indicating the inverse relationship between FAM99A levels and ribosomal gene expression. **(D)** Pearson Correlation analysis of FAM99A expression with 60S ribosomal protein genes in liver cancer. Scatter plots demonstrating strong negative correlations between FAM99A expression and essential 60S ribosomal subunit components including RPL13A (R = −0.29, p = 1.2e-11), RPL22 (R = −0.39, p = 3.4e-21), and RPL11 (R = −0.39, p = 2.2e-20) in TCGA liver cancer data. These consistent inverse correlations with both ribosomal subunit components support a model where FAM99A expression counteracts ribosome biogenesis and protein synthesis machinery.

These results provide direct functional evidence that FAM99A negatively regulates translation in liver cancer cells, consistent with our transcriptomic data showing downregulation of translation-associated pathways. The substantial reduction in protein synthesis suggests that FAM99A exerts a potent translational regulatory effect, which may contribute to its growth-inhibitory properties observed in the colony formation assays.

Collectively, these findings establish a functional link between FAM99A expression and global protein synthesis rates, revealing a mechanism through which this liver-specific lncRNA may exert its tumor-suppressive effects.

### FAM99A inversely correlates with ribosomal subunit gene expression

Analysis of TCGA liver cancer data revealed significant inverse correlations between FAM99A expression and key ribosomal protein genes (Figures 7C, 7D). We examined correlations with both small (40S) and large (60S) ribosomal subunit components. Among the 40S ribosomal proteins, FAM99A showed strong negative correlations with RPS6 (R = −0.34, p = 1.1e-15), RPS19 (R = −0.29, p = 9.1e-12), and RPS3 (R = −0.29, p = 1.3e-11) (Figure 7C). RPS6 is essential for translation initiation and serves as a key target of mTOR signaling (26), while RPS19 plays critical roles in pre-rRNA processing and ribosome assembly (27). Similarly, significant negative correlations were observed with 60S ribosomal proteins including RPL13A (R = −0.29, p = 1.2e-11), RPL22 (R = −0.39, p = 3.4e-21), and RPL11 (R = −0.39, p = 2.2e-20) (Figure 7D). RPL11 is particularly notable given its crucial role in ribosome biogenesis and p53 regulation through MDM2 interaction (28). These consistent negative correlations with both ribosomal subunit components support a model where FAM99A expression counteracts ribosome biogenesis and protein synthesis machinery at the transcriptional level. This relationship aligns with our observations of reduced global protein synthesis upon FAM99A overexpression and suggests coordination between transcriptional and post-transcriptional regulation of translation.

### TREX-MS identifies translation factors and components of ribosome biogenesis machinery as FAM99A protein binding partners

To identify proteins that directly interact with FAM99A, we employed Targeted RNase H-mediated Extraction of crosslinked RNA-binding proteins (TREX) in HepG2 cells, followed by liquid chromatography-tandem mass spectrometry (LC-MS/MS) and subsequent analysis. This approach allowed us to specifically capture proteins that physically associate with the FAM99A transcript under native cellular conditions. The TREX methodology identified a diverse array of proteins that interact with FAM99A (Figure 8A). The volcano plot showing differential protein enrichment between FAM99A and control samples revealed distinct clusters of significantly enriched proteins (blue and red dots, p < 0.05). Among the most significantly enriched proteins were several key components of the translation machinery, most notably eukaryotic translation initiation factors EIF4G3, EIF4G2, and EIF1AY, along with the RNA helicase DHX16 involved in ribosome biogenesis (Figure 8D).

**Figure 8:**
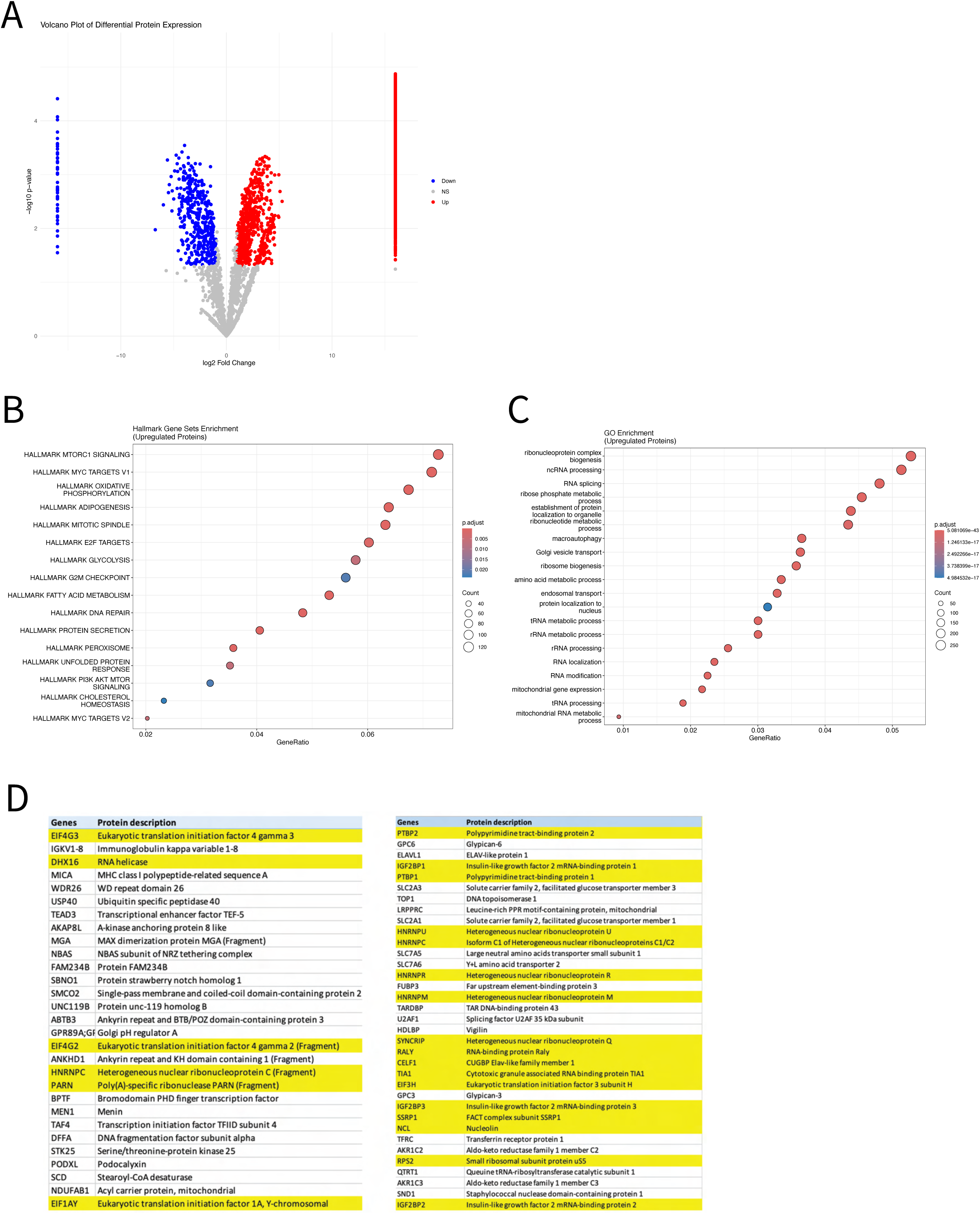
Identification of FAM99A protein-binding partners by TREX-MS. **(A)** Volcano plot showing differential enrichment of proteins captured in FAM99A versus control TREX-MS experiments. Significantly enriched proteins (p < 0.05) are highlighted in blue (downregulated) and red (upregulated). **(B)** Hallmark gene set enrichment analysis of upregulated FAM99A-binding proteins showing significant enrichment of pathways related to translation regulation and cellular metabolism. Circle size indicates gene count; color intensity represents adjusted p-value. **(C)** Gene Ontology enrichment analysis of FAM99A-binding proteins revealing predominant association with ribonucleoprotein complex biogenesis, rRNA processing, and ribosome biogenesis. **(D)** Top-ranked FAM99A-interacting proteins with example known translation and ribonuclear complex related hits highlighted in yellow.

Pathway enrichment analysis of FAM99A-interacting proteins using the Hallmark gene sets revealed significant enrichment of mTORC1 signaling, MYC targets, oxidative phosphorylation, and E2F targets (Figure 8B). The enrichment of these pathways aligns with the translation-related functions observed in our RNA-seq analysis. Particularly noteworthy was the strong enrichment of mTORC1 signaling, a master regulator of protein synthesis, and protein secretion pathways, highlighting FAM99A’s involvement in translation control. Gene Ontology (GO) analysis further substantiated the role of FAM99A in ribosome function and translation (Figure 8C). The most significantly enriched GO terms included ribonucleoprotein complex biogenesis, rRNA processing, and ribosome biogenesis. These findings are particularly consistent with our RNA-seq results, which showed minimal impact on mRNA transcript levels but significant downregulation of translation-related pathways upon FAM99A overexpression.

Our identification of translation initiation factors as FAM99A-binding partners provides a possible mechanistic explanation for the observed suppression of protein synthesis. EIF4G3 and EIF4G2 function as scaffolding proteins in the eIF4F complex that assembles translation machinery at mRNA 5’ caps. FAM99A’s interaction with these factors suggests it may directly inhibit translation initiation by interfering with eIF4F complex formation or function, thereby explaining how this lncRNA regulates protein synthesis in liver cells. Also of particular interest among the interacting proteins was DHX16, an ATP-dependent RNA helicase crucial for ribosome biogenesis and rRNA processing. This interaction, along with the enrichment of rRNA processing pathways, suggests that FAM99A may regulate translation not only at the initiation stage but also by influencing ribosomal RNA maturation and ribosome assembly. This finding is especially relevant considering that the liver is one of the most translationally active tissues in the body, with high demands for ribosome production.

Collectively, these results establish FAM99A as a regulatory lncRNA that directly interacts with key components of the translational machinery and ribosome biogenesis apparatus. These physical interactions provide a mechanistic foundation for the observed effects of FAM99A on protein synthesis and suggest a model where FAM99A serves as a molecular damper on translation in hepatocytes.

## DISCUSSION

Through integrative bioinformatic analysis, expression profiling, and functional studies, we have characterized FAM99A as a highly liver-enriched lncRNA that is consistently downregulated across liver malignancies and exerts growth-suppressive effects through the regulation of protein synthesis.

One of FAM99A’s most distinctive characteristics is its extreme tissue specificity. GTEx expression analysis demonstrated remarkably high expression in normal liver tissue (median 89.04 TPM), with negligible expression in all other tissue types examined. This pronounced hepatic enrichment suggests that FAM99A likely evolved to serve liver-specific functions, possibly related to the unique physiological demands of hepatocytes. FAM99A expression declined precipitously during conventional culture of primary human hepatocytes, with levels dropping to approximately 1% of initial values by 48 hours. This pattern parallels the well-documented phenomenon of hepatocyte dedifferentiation in standard two-dimensional culture (20,22,29). Interestingly, primary hepatocytes cultured under physioxic conditions (7% O₂) retained higher FAM99A expression compared to those in standard atmospheric oxygen, suggesting that more physiologically relevant culture conditions help preserve transcriptional programs driving FAM99A expression. Our isoform analysis identified FAM99A-203 as the predominant transcript in normal liver tissue. When our expression construct containing the longer FAM99A-201 cDNA was transfected into liver cells, we observed preferential processing to the FAM99A-203 form, indicating that hepatocyte splicing machinery favors this isoform even when presented with alternative templates.

Comparative analysis of FAM99A expression across liver malignancies revealed systematic downregulation in all tumor types examined, with the degree of suppression varying by cancer subtype. This consistent pattern of downregulation suggests strong selective pressure against FAM99A expression during hepatic malignant transformation, and/or loss of FAM99A as a liver-specific transcript during dedifferentiation from a native hepatic state. The clinical significance of FAM99A expression was demonstrated through survival analysis of HCC patients. Those with higher FAM99A levels exhibited significantly better overall survival (HR = 0.58, 95% CI: 0.41-0.82, p = 0.0019), with the survival advantage becoming particularly evident beyond 40 months post-diagnosis. This long-term survival benefit suggests that FAM99A may influence fundamental aspects of disease progression rather than merely correlating with other prognostic factors.

Our functional studies provided direct evidence for the tumor-suppressive capacity of FAM99A. Transient overexpression in liver cancer cell lines reduced anchorage-independent growth in soft agar, a hallmark of malignant transformation. The most mechanistically informative finding was the discovery that FAM99A negatively regulates global protein synthesis rates. Transcriptome analysis in both Huh7 cells and primary human hepatocytes indicated the repression of translation-associated pathways upon FAM99A overexpression. Gene Set Enrichment Analysis demonstrated significant negative enrichment in multiple translation-related processes, including “Translation,” “Eukaryotic Translation Elongation,” “Formation of a pool of free 40S subunits,” and “Ribosome biogenesis.” The consistent suppression of translation-related pathways across cell models establishes translational regulation as a primary function of FAM99A in both normal and malignant liver cells. Our OPP incorporation assay demonstrated that FAM99A overexpression substantially reduced translation in both Hep3B and Huh7 cells. This dramatic effect on protein synthesis provides a plausible mechanism for FAM99A’s growth-suppressive properties, as rapidly proliferating cancer cells are particularly dependent on elevated translation rates to support increased biosynthetic demands.

TREX-MS analysis identified multiple components of the translation machinery as direct binding partners of FAM99A, including eukaryotic translation initiation factors EIF4G3, EIF4G2, and EIF1AY. This interaction provides a mechanistic explanation for our observations that FAM99A overexpression suppresses global protein synthesis rates. EIF4G3 and EIF4G2 serve as scaffolding proteins in the eIF4F complex, which coordinates the assembly of translation initiation machinery at mRNA 5’ caps. Their interaction with FAM99A suggests that this lncRNA may directly modulate translation initiation by interfering with eIF4F complex formation or function. Also of interest among the interacting proteins was DHX16, an ATP-dependent RNA helicase crucial for ribosome biogenesis and rRNA processing. This interaction, along with the enrichment of rRNA processing pathways, suggests that FAM99A may regulate translation not only at the initiation stage but also by influencing ribosomal RNA maturation and ribosome assembly.

FAM99A bears functional similarities to its paralog FAM99B, which has been characterized as a tumor suppressor in HCC (16,17). Both genes show liver-enriched expression and downregulation in liver cancers. FAM99B regulates ribosome biogenesis by interacting with the RNA helicase DDX21 and promoting its proteolytic degradation (17). This functional convergence on protein synthesis machinery suggests that the FAM99A/B gene pair may have evolved specialized roles in controlling translation in hepatocytes, potentially reflecting the extraordinary protein synthesis demands of the liver.

The liver’s unique physiological context provides important perspective for interpreting FAM99A’s role in translational regulation. Hepatocytes are among the most translationally active cells in the body, producing large quantities of plasma proteins, metabolic enzymes, and detoxification factors (30,31). This high secretory load requires precise regulation of protein synthesis to balance productive translation with cellular homeostasis. As a liver-specific negative regulator of translation, FAM99A may help calibrate protein synthesis rates to prevent excessive translational burden on hepatocytes.

Our observation that HBV reduces FAM99A expression in hepatocytes reveals a potential mechanism by which this oncogenic virus may promote carcinogenesis. Since FAM99A suppresses protein synthesis, its HBV-mediated downregulation could enhance translation rates in infected cells, simultaneously facilitating viral replication and cellular transformation (32–34). This finding provides a novel link between HBV infection, translational control, and hepatocarcinogenesis that warrants further investigation.

Future research should address several key aspects of FAM99A biology. First, detailed investigation of the precise mechanism by which FAM99A regulates translation is warranted. Ribosome profiling experiments would enable comprehensive assessment of FAM99A’s impact on translational efficiency across the transcriptome, revealing whether it exerts global effects or selectively regulates specific mRNA subsets. Second, improved model systems would facilitate more rigorous functional characterization. Three-dimensional culture systems such as liver spheroids or organoids would better recapitulate the native hepatic environment and could preserve FAM99A expression patterns lost in conventional 2D culture. Patient-derived organoids would provide valuable platforms for studying FAM99A regulation in the context of viral infection and tumorigenesis. Finally, exploration of FAM99A as a potential therapeutic target or biomarker holds promise. The consistent downregulation across liver cancer subtypes suggests utility in diagnostic applications, while its correlation with tumor stage and survival indicates prognostic value. Strategies to restore FAM99A function in HCC warrant exploration, including direct delivery of synthetic FAM99A RNA using hepatocyte-targeting nanoparticles or development of small molecules that mimic FAM99A’s translational regulatory effects. We also show evidence that the presence of HBV may downregulate FAM99A expression. When paired with the appearance of ‘viral mRNA translation’ as a significantly enriched ReactomePA pathway in the Huh7 RNA-seq analysis of FAM99A overexpression, these data imply a potential interplay between HBV and FAM99A warranting future investigation. 3’ and 5’RACE techniques followed by sequencing can also assist in experimentally verifying any novel FAM99A isoforms and improve current annotations.

In conclusion, we have identified FAM99A as a liver-specific lncRNA that functions as a negative regulator of protein synthesis and exhibits tumor-suppressive properties in hepatocellular carcinoma cell lines. This work establishes a foundation for understanding FAM99A’s role in liver biology and cancer, opening numerous avenues for further investigation with refined experimental approaches and expanded model systems.

## MATERIALS AND METHODS

### Data Sources

RNA sequencing data from multiple sources were integrated to identify and characterize liver-specific long non-coding RNAs. The primary datasets included: (1) The Cancer Genome Atlas Liver Hepatocellular Carcinoma (TCGA-LIHC) cohort for tumor versus normal liver differential expression analysis; (2) Genotype-Tissue Expression (GTEx v10) dataset for tissue-specificity analysis across 54 human tissues; and (3) RNA-seq data from liver cancer cell lines and primary human hepatocytes (PHHs). TCGA-LIHC raw count matrices were retrieved through the TCGAbiolinks R package and GTEx v10 median TPM expression data were obtained from the GTEx portal (GTEx_Analysis_2022-06-06_v10_RNASeQCv2.4.2_gene_median_tpm.gct). Additional RNA-seq data for different liver cancer subtypes (HCC, HPBL, CHOL, and FLC) were obtained from publicly available datasets in the Gene Expression Omnibus (GEO) and processed through the pipeline described below, selecting only samples with minimum sequencing depth of 20 million reads generated from total RNA extractions. Detailed metadata for these samples are provided in Supplementary Table S1. For experimental samples (liver cancer cell lines and primary human hepatocytes), RNA sequencing was performed by Azenta Genewiz, and the raw sequencing files were processed through the bioinformatics pipeline described below.

### RNA-seq Data Processing

Raw RNA-seq data were processed using stratoolkit3 and the nf-core/rnaseq pipeline (version 3.11.2) with Nextflow (version 22.10.1) and Singularity container engine. Alignment used the human reference genome GRCh38.p13 with GENCODE v43 annotations. The pipeline incorporated FastQC for quality control, Trim Galore for adapter trimming, STAR for genome alignment, Salmon for transcript quantification, and RSeQC for aligned data quality control. STAR-aligned BAM files were processed with StringTie (version 2.2.1), and count matrices were generated for downstream analysis in R. All processing was performed on a high-performance computing cluster.

### Differential Expression and Correlation Analyses

Differential expression analysis was performed using DESeq2 to identify genes with significant expression changes between different sample groups. For the identification of liver-specific lncRNAs, the TCGA-LIHC cohort was analyzed to compare tumor versus normal liver tissue samples. Raw count matrices were filtered to remove low-count genes (fewer than 10 counts across all samples). Size factors were estimated using the median ratio method, and dispersion estimates were obtained using a parametric fit. The Wald test was used for statistical inference with Benjamini-Hochberg correction for multiple testing. Genes were considered differentially expressed at an adjusted p-value < 0.05 and |log₂ fold change| ≥ 1.

For the cross-cancer-type analysis comparing normal liver, HCC, HPBL, FLC, and CHOL samples, batch effects were addressed using ComBat-seq prior to differential expression analysis. Following batch correction, the data were transformed using variance-stabilizing transformation (VST) to account for the mean-variance relationship inherent in RNA-seq data. Principal component analysis (PCA) and hierarchical clustering were performed on VST-transformed data to assess global relationships between samples.

Statistical significance for pairwise comparisons between individual cancer types and normal liver was assessed using either Student’s t-test (for normally distributed data) or Wilcoxon rank-sum test (for non-normal distributions), depending on data distribution as determined by Shapiro-Wilk test. P-values were adjusted for multiple testing using the Benjamini-Hochberg method, with significance defined as adjusted p < 0.05.

For visualization of differential expression across TCGA cohorts, the GEPIA2 web portal was utilized to generate expression plots for both TCGA-LIHC and TCGA-CHOL datasets. Similarly, the GEPIA2 web portal was utilized for custom Pearson correlation analyses between FAM99A expression and ribosomal subunit gene expression using LIHC Tumor, LIHC Normal, and Liver expression datasets.

### Identification of Liver-Specific lncRNAs

To identify liver-specific lncRNAs, expression patterns across multiple tissue types were analyzed using the GTEx v10 dataset. LncRNAs were defined as transcripts annotated as “lincRNA,” “antisense,” “processed_transcript,” “sense_intronic,” “sense_overlapping,” “3prime_overlapping_ncRNA,” “long_non_coding,” “bidirectional_promoter_lncRNA,” “macro_lncRNA,” or “lncRNA” in Ensembl BioMart (release 113) or NCBI RefSeq annotations. Liver specificity was determined using two criteria: (1) high expression in liver (≥10 TPM), and (2) at least 5-fold higher expression in liver compared to any other tissue. This stringent approach identified 18 liver-specific lncRNAs, which were then intersected with differentially expressed lncRNAs from the TCGA-LIHC analysis to prioritize candidates for functional characterization. The GTEx dataset was utilized to create a tissue expression heatmap, where expression values across 54 human tissues were normalized and visualized. RNA-seq read coverage from the TCGA-LIHC dataset was examined to validate the expression patterns of candidate liver-specific lncRNAs in both tumor and adjacent normal samples.

### Survival Analysis and Clinical Correlation

Kaplan-Meier survival analysis was performed to evaluate the clinical significance of FAM99A in HCC patients using the Kaplan-Meier Liver Cancer RNA-seq web portal (https://www.kmplot.com/analysis/) with auto-select of best cutoff percentile and no restriction of subtypes (n=364).

### Transcript Isoform Analysis

For isoform-specific expression analysis across various liver tissues, transcript count matrices obtained as described above were filtered to FAM99A transcripts, and expression values were normalized using log2(counts + 1) transformation. Transcript usage was compared across normal liver, HCC, FLC, HPBL, and CHOL samples, with expression patterns visualized using custom scripts implemented in R. Visual representations were generated in the form of split violin plots and heatmaps to illustrate isoform-specific expression distributions across tissue types.

Splice junction analysis was performed using the Integrative Genomics Viewer (IGV v2.16.2) with a normal liver biopsy sample (SRR8601557) BAM file generated through the pipeline described above and the human GRCh38 reference genome. RNA-seq read coverage was examined to validate expression patterns of FAM99A transcript variants in both tumor and adjacent normal samples. Custom sashimi plots were generated to visualize splice junctions and exon usage across samples. The plots were configured to display read coverage along with arcs representing splice junctions, with the thickness of the arcs proportional to the number of reads supporting each junction. These visualizations enabled the identification of predominant transcript isoforms and alternative splicing events in FAM99A across different tissue samples.

Experimental validation of FAM99A transcript isoforms was conducted using RT-PCR. Total RNA was extracted from cell samples using TRIzol reagent according to the manufacturer’s protocol. RNA integrity was qualitatively assessed using 1% agarose gel electrophoresis to visualize sharp rRNA bands, and both quality and quantity were evaluated using a NanoDrop spectrophotometer. Complementary DNA (cDNA) was synthesized using NEB M-MuLV reverse transcriptase with modifications to the manufacturer’s protocol.

Briefly, a reaction mixture containing 2 μL of 10× M-MuLV RT Buffer, 3 μL of 25 μM random hexamers, 1 μL of 100 μM oligo(dT), 2 μg of RNA, and nuclease-free H₂O to a volume of 10 μL was prepared. The mixture was incubated at 75°C for 10 minutes, followed by 4°C for 5 minutes. Subsequently, 1 μL of M-MuLV RT (200 U/μL) for +RT samples or 1 μL of nuclease-free H₂O for -RT controls, 1 μL of 10 mM dNTP mix, 0.5 μL of RNase inhibitor (40 U/μL), and nuclease-free H₂O to a final volume of 20 μL were added. The reaction was incubated at 25°C for 5 minutes, 37°C for 1 hour, 95°C for 5 minutes, and finally held at 4°C. The synthesized cDNA was either used immediately or stored at −20°C for later use. PCR amplification was performed using NEB Taq 2× Master Mix according to the manufacturer’s protocol in a 25 μL reaction format, with 2.5 μL of synthesized cDNA per reaction. Primers were designed to target regions that would yield distinct amplicon sizes for different annotated FAM99A transcript variants (Ensembl release 112, May 2024): FAM99A-201 (903 bp), FAM99A-202 (1061 bp), and FAM99A-203 (204 bp), determined using the USCS In-Silico PCR browser (Human Genome, Assembly Dec. 2013 [GRCh38/gh38], Target GENCODE genes). The primer sequences used were:

FP (5’->3’): ATGGAGAAGGCTCCCTGT
RP (5’->3’): ACATAGGATTCCCACAGTG

PCR products were resolved on 1% agarose gels, and band sizes were compared to predicted amplicon sizes to identify the predominant transcript isoforms expressed in different cell types. This approach provided experimental validation of the isoform usage patterns observed in the RNA-seq data analysis.

### Functional Enrichment Analysis

Gene Set Enrichment Analysis (GSEA) was performed to identify coordinated changes in gene expression across established biological pathways. The fgsea R package (version 1.30.0) was employed with default parameters. Ranked gene lists were generated by ordering genes based on the signed log2 fold-change values from differential expression analyses, providing directionality (up/down-regulation) of expression changes. Normalized enrichment scores (NES) and adjusted p-values (using Benjamini-Hochberg correction) were calculated to assess pathway significance (p-adjusted < 0.05).

Multiple gene set collections were interrogated to ensure comprehensive evaluation of enriched biological processes. These included Gene Ontology (GO) biological process, molecular function, and cellular component terms; Kyoto Encyclopedia of Genes and Genomes (KEGG) pathways; and MSigDB Hallmark gene sets. For GO-based enrichment, both standard over-representation analysis and GSEA-style rank-based methods were applied using the clusterProfiler R package (version 4.12.6). KEGG pathway enrichment was conducted for protein-coding genes with mapIds to enable retrieval of ENTREZID identifiers. The ReactomePA package was utilized to investigate Reactome pathway enrichment, providing additional coverage of canonical signaling cascades.

To identify FAM99A-specific effects in primary human hepatocytes (PHHs), a strategic analytical approach was implemented to control for artifacts introduced by adenoviral transduction. PHHs from three independent donors were transduced with adenoviral vectors expressing either FAM99A (pAV-FAM99A) or enhanced green fluorescent protein (pAV-GFP) as a control, or left untransduced. RNA-seq was performed 72 hours post-transduction. A differential expression comparison between untransduced and pAV-GFP samples was conducted to identify gene programs altered by viral transduction and/or GFP alone, establishing a transduction/GFP-effect signature. Subsequently, all gene sets significantly enriched (p-adjusted < 0.05) in the pAV-GFP versus untransduced comparison were excluded from the pAV-FAM99A versus pAV-GFP comparison to ensure that reported pathway alterations were specifically attributable to FAM99A expression rather than general transduction effects. Additional conservative filtering was applied to classify genes based on expression patterns; genes showing concordant changes in both pAV-GFP and pAV-FAM99A were classified as “Transduction Effect,” while genes showing significant changes only in pAV-FAM99A samples or exhibiting significantly stronger magnitude changes compared to pAV-GFP were classified as “FAM99A-Specific”.

### TREX-MS FAM99A-Interacting Protein Analysis

RNA-protein interactions were captured using the Targeted RNase H-mediated Extraction of crosslinked RNA-binding proteins (TREX) methodology as described by Dodel et al. (35). HepG2 cells (2 × 10^7) were subjected to UV-C crosslinking to covalently stabilize in vivo RNA-protein interactions. Crosslinked cells were lysed in TRIzol, followed by phase separation with chloroform. This separation technique resulted in free RNA partitioning to the aqueous phase and free proteins to the organic phase, while RNA-protein adducts accumulated at the interface due to their insolubility. The interphase fraction was isolated and washed repeatedly with TE buffer until TRIzol dye was completely removed. Sequential dissolution of the interphase material was performed using TE buffer containing increasing concentrations of SDS (0.1% and 0.5%), with centrifugation steps between each extraction to collect supernatants. RNA-protein complexes were precipitated from the combined supernatants by addition of sodium chloride (5M), glycogen, and isopropanol, followed by centrifugation at 18,000g at 4°C. The resulting pellet was washed with 70% ethanol, consolidated, and hydrated in 360μL HyPure water on ice.

Contaminating DNA was eliminated through digestion with TURBO DNase (100μL) in the presence of RNase inhibitor (80U) for 50 minutes at 37°C with agitation. Following precipitation and washing, the sample was resuspended in hybridization buffer containing antisense DNA oligonucleotides specifically designed to target FAM99A transcript (Table 2). DNA-RNA hybridization was achieved through controlled temperature reduction from 95°C to 50°C. Thermostable RNase H was then applied to digest the DNA-RNA hybrids, resulting in the release of FAM99A-associated proteins. The liberated RNA-binding proteins were extracted through TRIzol LS treatment (900μL) followed by chloroform phase separation (200μL) and overnight acetone precipitation at −-20°C. The protein precipitate was washed with 80% acetone, air-dried, and resuspended in HyPure water before submission for liquid chromatography-tandem mass spectrometry (LC-MS/MS) analysis. Protein identification and quantification data were analyzed using clusterProfiler to determine enriched pathways associated with FAM99A-binding proteins.

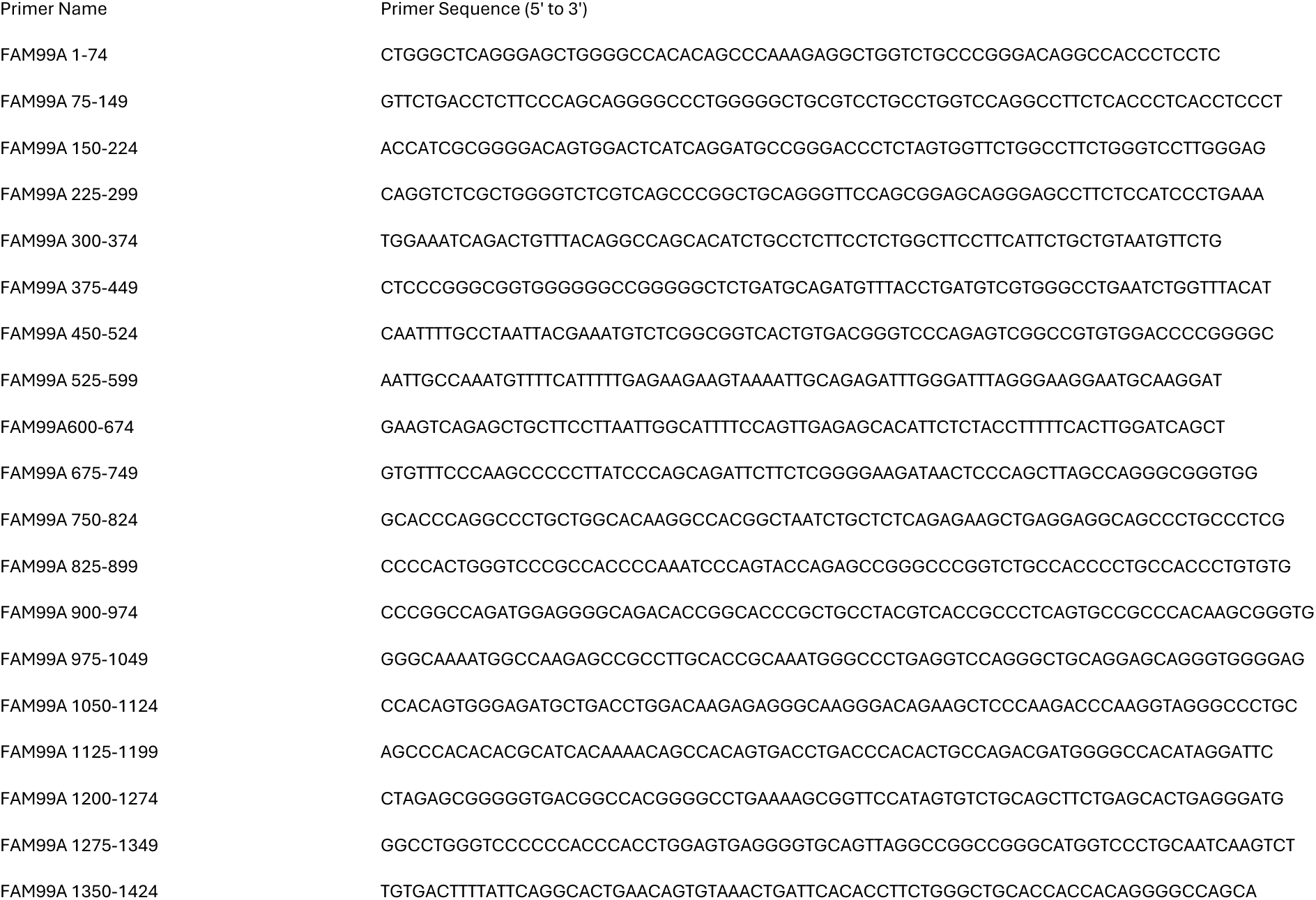

Pathway enrichment analysis of FAM99A-binding proteins was conducted using MSigDB Hallmark gene sets, with significance of enrichment quantified by p-values. Gene Ontology (GO) analysis of FAM99A-interacting proteins was performed to identify overrepresented biological processes, molecular functions, and cellular components. This integrative approach enabled the characterization of FAM99A’s molecular interaction network and provided insights into its functional role in translation regulation.

### Statistical Analysis

All statistical analyses were performed in R (version 4.4.0) or GraphPad Prism 10. For comparisons between two groups where each experimental replicate was normalized to its matched control (set to 1), one-sample t-tests were applied to determine if the experimental group means differed significantly from the theoretical value of 1. For standard two-group comparisons without normalization, Student’s t-test or Wilcoxon rank-sum test was used depending on data normality. For multiple group comparisons, ANOVA or Kruskal-Wallis tests were applied, followed by post-hoc tests with Benjamini-Hochberg correction. Statistical significance was defined as p < 0.05 unless otherwise specified.

### Cell Lines and Culture Conditions

Liver cancer cell lines (HepG2, Huh7, and Hep3B) were maintained in Minimum Essential Medium (MEM) supplemented with 10% fetal bovine serum (FBS), 1% sodium pyruvate, 1% L-glutamine, 1% non-essential amino acids, and 1% penicillin-streptomycin. Cells were cultured at 37°C in a humidified atmosphere containing 5% CO₂. Primary human hepatocytes (PHHs) were isolated from donor liver tissues from the University of Pittsburgh Human Liver Tissue and Hepatocyte Research Resource (HLTHRR). Upon arrival, PHHs were maintained in Williams’ E Medium (WEM) supplemented with 1% sodium pyruvate, 1% L-glutamine, 0.4% Insulin-Transferrin-Selenium (ITS), 0.1% hydrocortisone, 0.02% human epidermal growth factor (EGF), and 0.06% gentamicin, and plated on 5x-collagen-coated polystyrene.

### Transfections and Transductions

Transient transfections were performed in collagen-coated 6-well plates using GeminiBio Continuum transfection reagent according to the manufacturer’s protocol. Briefly, optimal Cotninuum:DNA ratios were first assessed per cell type using pGEM-GFP transfections and fluorescence microscopy. Once determined, 1 μg of plasmid DNA (pcDNA3.1-FAM99A or empty vector control pcDNA3.1ΔNE) was mixed with the transfection reagent at the optimal ratio and added to cells at 70-80% confluence.

For adenoviral transduction, primary human hepatocytes were infected with adenoviral vectors pAV-FAM99A (pAV[ncRNA]-CMV>hFAM99A[NR_026643.1] designed with VectorBuilder) or pAV-EGFP 3 hours after plating at a multiplicity of infection (MOI) of 1 per VectorBuilder instructions. Transduction efficiency was monitored by fluorescence microscopy of GFP expression.

### RNA Extraction and RT-qPCR

Total RNA was extracted using TRIzol reagent, quantified by NanoDrop spectrophotometry, and analyzed by RT-qPCR (qScript One-Step SYBR Green kit) using 100 ng RNA per reaction on a Bio-Rad CFX96 system. FAM99A primers (forward: 5’-CACGACATCAGGTCAGCATCT-3’; reverse: 5’-AAAAGCGGTTCCATAGTGTC-3’) and GAPDH primers (forward: 5’-TCGGAGTCAACGGATTTGGT-3’; reverse: 5’-TTCCCGTTCTCAGCCTTGAC-3’) were used. Expression levels were calculated using the 2^(-ΔΔCt) method with GAPDH normalization.

### Anchorage-Independent Growth Assay

Soft agar colony formation assays were performed to assess anchorage-independent growth capabilities. Briefly, a base layer of 2mL 0.5% low-melting temperature agarose in complete growth medium was prepared in 6-well plates and allowed to solidify. In parallel, 24-hour post-transfection cells were trypsinized, pelleted and resuspended in fresh media, and counted. The appropriate amount of cells was resuspended in a top layer of 1mL 0.3% low-melting temperature agarose cooled to 40°C in complete growth medium to obtain a density of 10,000 cells per well. After solidification of the top layer, 500 uL of complete growth medium was added to each well and refreshed every 2 days. Colonies were allowed to form for 14 days, after which they were fixed and stained overnight with 1mL/well of 0.05% p-iodonitrotetrazolium violet prepared in a 50/50 solution of deionized water and methanol, imaged using a EVOS FL Auto system, and counted using a custom Python application.

### Protein Synthesis Assay

Global protein synthesis rates were measured using the Protein Synthesis Assay Kit (Cayman Chemical, Item No. 601100) according to the manufacturer’s protocol using black, clear-bottom 96-well plates. This assay utilized O-propargyl-puromycin (OPP), which was incorporated into nascent polypeptide chains and subsequently detected via copper-catalyzed click chemistry with a fluorescent azide. Briefly, cells were incubated with OPP for 1.5 hours, fixed, and labeled with 5-FAM-azide. Fluorescence intensity was quantified using a BioTek Synergy 2 Hybrid Multi-Detection Microplate Reader (bottom-read, auto-gain) and qualitatively assessed via microscopy with a EVOS FL Auto system. Cells were counterstained with DAPI to visualize and count nuclei, and fluorescence intensity was normalized to cell number to obtain per-cell protein synthesis rates.

